# Mechanisms and implications of high depolarization baseline offsets in conductance-based neuronal models

**DOI:** 10.1101/2024.01.11.575308

**Authors:** Anal Kumar, Anzal K. Shahul, Upinder S. Bhalla

## Abstract

Somatic step-current injection is commonly used to characterize the electrophysiological properties of neurons. Many neuronal types show a large depolarization baseline offset (DBLO), which is defined as the positive difference between the minimum membrane potential during action potential trains and resting. We used stochastic parameter search in experimentally constrained conductance-based models to show that four key factors together account for high DBLO: Liquid Junction Potential correction, high backpropagating passive charges during the repolarization phase of an action potential, fast potassium delayed rectifier kinetics, and appropriate transient sodium current kinetics. Several plausible mechanisms for DBLO, such as Ohmic depolarization due to current input or low-pass filtering by the membrane, fail to explain the effect, and many published conductance-based models do not correctly manifest high DBLO. Finally, physiological levels of DBLO constrain ion channel levels and kinetics, and are linked to cellular processes such as bistable firing, spikelets, and calcium influx.

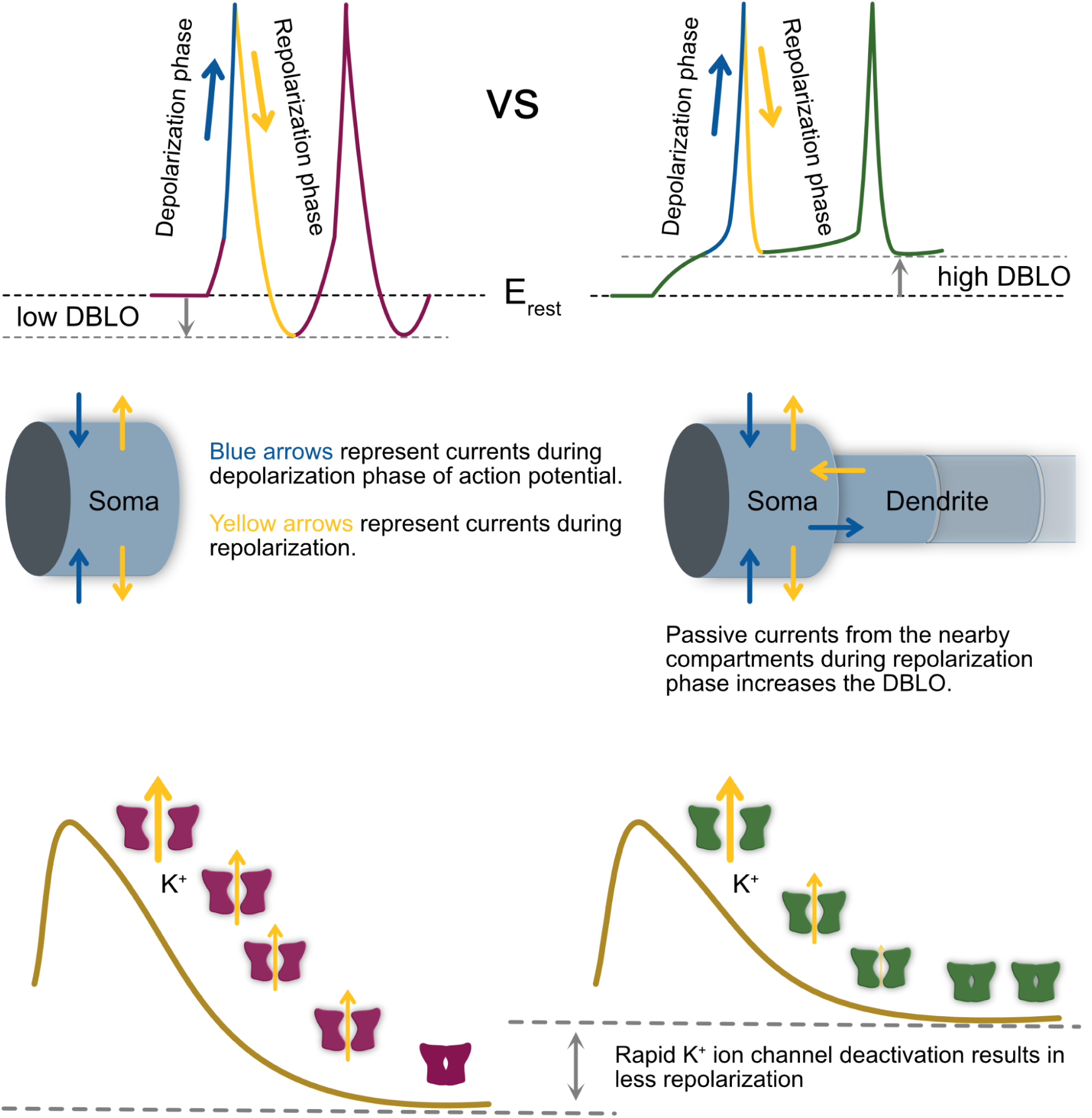

## Introduction

Accurate neuronal models have become essential tools in experimental and computational neuroscience. Key details of action potential (AP) shape provided early clues about the existence and roles of various ion channels in different cell types. For example, the afterhyperpolarization (AHP) following an AP in the sinoatrial (SA) node of the heart led to the discovery of “funny” (HCN) channels (H. F. Brown et al., 1979). Although extant neuronal models have been instrumental in providing insights into the biophysics of neurons, e.g., in understanding back-propagating APs (M. Migliore et al., 1999) and calcium spikes (Miyasho et al., 2001), some key details have remained understudied. One such detail is the high Depolarization BaseLine Offset (DBLO) seen in somatic recordings when long pulses of current are injected into a neuron (Bianchi et al., 2022). High DBLO can be seen in current-clamp recordings of almost all pyramidal neuron subtypes: hippocampal CA1 (Dwivedi et al., 2019), CA3 (Hunt et al., 2018), cortical L2/3 (Moradi Chameh et al., 2021) and L5 (Kim et al., 2015; Moberg & Takahashi, 2022). Many other neurons, including thalamocortical neurons (Studtmann et al., 2023) and olfactory bulb M/T cells (Fadool et al., 2011), also exhibit high DBLO. Neurons with low DBLO include fast-firing basket cells (Pawelzik et al., 2003; Szabó et al., 2010). Notably, the original Hodgkin-Huxley (HH) squid recordings displayed low DBLO (Hodgkin & Huxley, 1952), and this has tended to dominate illustrations of neuronal firing (Kandel et al., 2021). Since high DBLO is also seen in perforated patches (Hess et al., 2021; Staff et al., 2000) and opsin-induced stimulations (Alejandre-García et al., 2022), it is unlikely to be a method-specific phenomenon.

Several studies have hinted at the role of morphology in the phenomenon of high DBLO. Pinching or cutting off dendrites leads to the disappearance of high DBLO (Bekkers & Häusser, 2007). Dissociated CA1 pyramidal neurons, which are essentially neurons without the dendritic and axonal architecture, show low DBLO (Liu & Bean, 2014). Furthermore, a modeling study (Hendrickson et al., 2011) showed that high axial currents from dendrites resulted in a reduction in the amplitude of fast afterhyperpolarization (fAHP), which is a proxy of high DBLO in these spontaneously firing neuron models. It can be argued that a sustained depolarizing current can give rise to high DBLO. However, blocking persistent sodium channels (Na_P) does not lead to low DBLO (Müller et al., 2018). Calcium channels also do not seem to have a significant effect on DBLO, as blocking Ca^2+^ channels or changing Ca^2+^ levels had a limited impact on fAHP and, thus, DBLO (Sahu et al., 2017, 2019). Further, blocking R-type Ca^2+^ channels also did not have a significant effect (Metz et al., 2005).

Modelers have largely ignored the phenomenon of high DBLO. As we analyze below, several models of CA1 pyramidal neurons on ModelDB do not show the experimentally expected level of DBLO in current clamp recordings. Recently, (Bianchi et al., 2022)) addressed this issue and suggested that the high Depolarization Baseline (DBL) may be due to an injection-current dependent shift in reversal potential on ion channels as well as their kinetics (Bianchi et al., 2022). The mechanistic basis for such a proposed current-dependent shift is uncertain.

With current morphological (Ascoli et al., 2007; Gouwens et al., 2018) and ion channel (Podlaski et al., 2017; Ranjan et al., 2011) databases, easy access to powerful computers and simulators (Alam et al., 2022; Carnevale & Hines, 2006; Eppler et al., 2009; Goodman & Brette, 2008; Ray & Bhalla, 2008), it is easy to build complex models which may in some cases fit well to experimental data without providing insights into neuronal phenomenon. This is pronounced if a phenomenon is a composite outcome of multiple mechanisms, which, as we find, is the case for DBLO. Thus, we found it valuable to work through minimal models to pin down individual mechanistic contributions. To do this, we systematically checked the contributions of various factors in conductance-based models of CA1 pyramidal neurons that affect DBLO and contribute to high DBLO.

In the current study, we obtained electrophysiological data from mouse CA1 pyramidal neurons for passive properties, frequency response, and DBLO with current clamp recordings. We used this data to build models and to explore factors that affect DBLO. We found that multiple factors contribute to DBLO, including passive properties, morphology, and channel kinetics. Although the factors mentioned above were not strong enough to produce a large number of high DBLO models independently, clubbing them together produced many models with high DBLO. Furthermore, high DBLO plays a nuanced role in certain signature properties of neuronal firing, such as firing bistability due to persistent sodium channels. We also show that DBLO levels in the models are correlated with internal calcium concentration, and the minimum gap junction conductance needed for the induction of an AP in gap junction paired neurons.

## Results

### CA1 pyramidal neurons exhibit high DBLO

To characterize DBLO in CA1 pyramidal neurons, we recorded membrane potential from mouse hippocampal CA1 pyramidal neurons (N=13) while injecting a series of step currents for 500 ms. Fig 1A and Fig 1B show representative recordings from the soma of a neuron when 150 pA and 300 pA, respectively, were injected into the soma. We calculated the DBLO of these recordings, which is denoted by the arrows between the horizontal dashed and dotted lines in Figs. 1A and 1B. The DBL is defined as the mean of the minimum membrane potential between two consecutive spikes, and the DBLO is defined as the difference between DBL and the resting membrane potential (E_rest_) of the neuron. At 150 pA, the mean DBLO at 150 pA current injection across all the 13 neurons was 18.8 ± 4 mV (range of 11.6 mV to 27.6 mV). The 10 to 90 percentile range of DBLO at 150 pA current injection was 14.3 mV to 23.6 mV. For our study, we chose this as our convention for labeling a DBLO as “high” (14.3 to 23.6 mV). DBLO values between 10 and 14.3 mV were classified as “moderate,” while those below 10 mV were considered “low”.

**Fig. 1.**
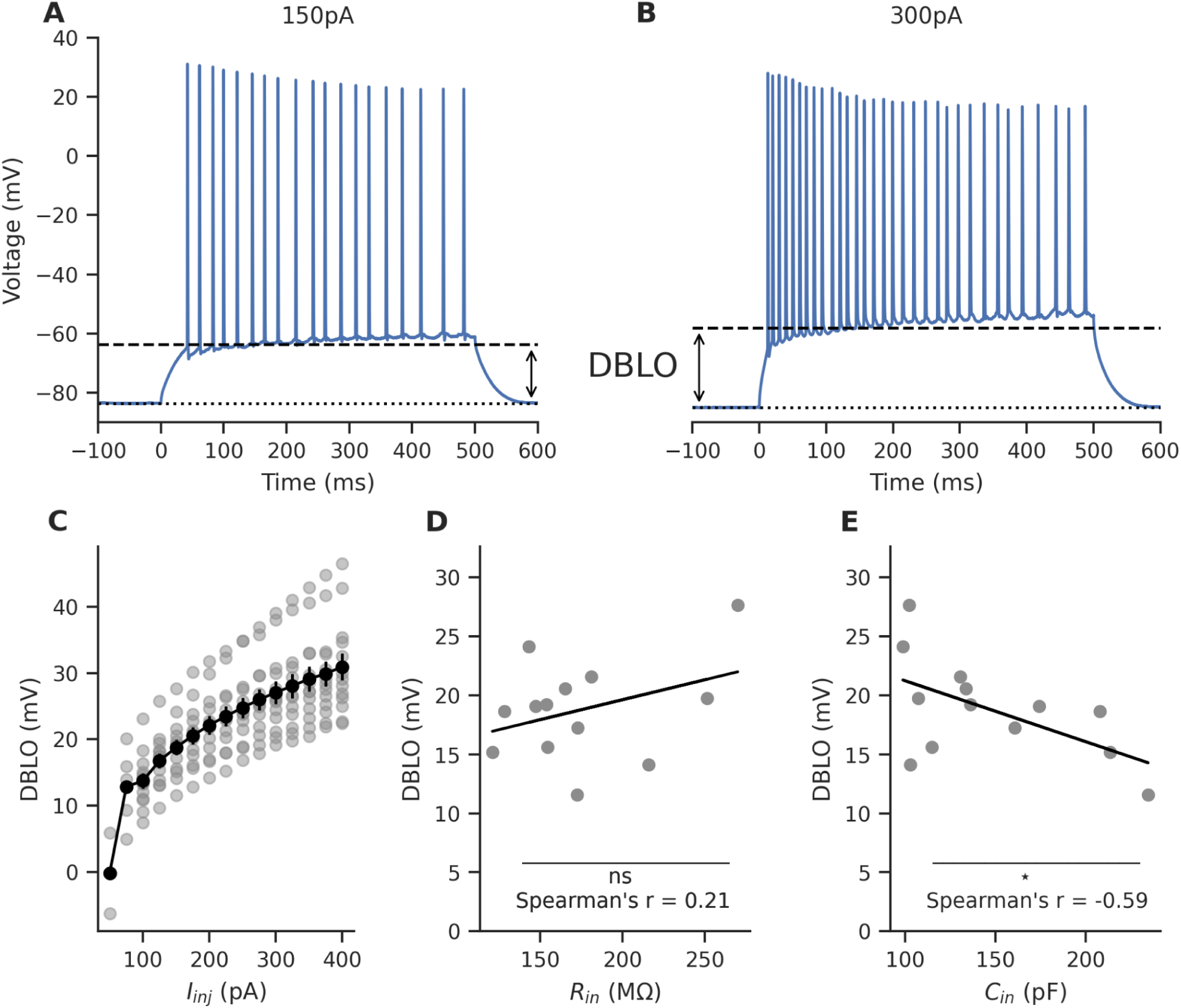
Experimentally observed DBLO features. **(A)** A representative whole-cell recording from a mouse CA1 pyramidal neuron showing a DBLO of 19.7 mV. 150 pA current was injected into the soma of the neuron for 0.5 s, starting at t = 0 s. The horizontal dashed and solid lines represent the DBL and E_rest_ of the neuron, respectively. The potential difference between these two represents the DBLO. **(B)** Same as A but for 300 pA current injection. The DBLO here is 26.8 mV. **(C)** The DBLO increases with the amplitude of the injected current. Data from 13 recorded neurons are shown in gray. The black dots represent the mean DBLO at each injected current level. The error bars represent one standard error of the mean (sem). **(D)** DBLO shows a non-significant correlation (Spearman’s correlation coefficient r = 0.21, p-value = 0.49) with the input resistance (R_in_) of the neuron. **(E)** DBLO shows a moderate negative correlation (spearman’s correlation coefficient r = −0.58, p-value = 0.035) with the total cell capacitance (C_in_) of the neuron.

The DBLO of neurons increased with the amount of current injected (Fig. 1C). Although Ohm’s law (ΔV = I*R) predicts this, it could not explain the absence of correlation between the input resistance (R_in_) and the DBLO (Fig. 1D, Spearman’s correlation coefficient r = 0.21, p-value = 0.49). On the other hand, the neuron’s total capacitance was found to be moderately inversely correlated with DBLO (Fig. 1E, Spearman’s correlation coefficient r = −0.59, p-value = 0.035). Thus, in this first section, we experimentally characterized DBLO from mouse CA1 pyramidal neurons.

### Numerous conductance-based models of CA1 pyramidal neurons have low DBLO

To examine how DBLO has been addressed in the simulation literature, we looked at some state-of-the-art CA1 pyramidal neuron conductance-based models with detailed morphology. We first calculated the DBLO in a set of pyramidal neuron models from ModelDB (McDougal et al., 2017) and the Allen Cell Types Database (http://celltypes.brain-map.org/). The original models were implemented in the NEURON (Carnevale & Hines, 2006). Hence, we used NEURON for these simulations. Since these models were made with information from cells of different animals and brain regions, they may show differences in intrinsic properties like input resistance and resting membrane potential. To account for these variations in the intrinsic properties of each model, we injected a current adjusted to produce the same number of spikes—19—in 500 ms. This target of 19 spikes was chosen to match the representative recording shown in Fig. 1A, where a 150 pA current injection yielded 19 spikes. This approach also avoided any potential effects that may have arisen from differences in the firing rates and excitabilities of the models.

The model of rat CA1 pyramidal neuron from (R. Migliore et al., 2018) showed a DBLO of only 1 mV (Fig. 2A). As we discuss in detail in the next section, liquid junction potential (LJP) correction is vital for building models with high DBLO. Since this model did not correct their experimental recordings for LJP, we compensated for the LJP in the models by decreasing the leak reversal potential (E_m_) of the model such that the resulting E_rest_ of the model was 15 mV lower than the E_rest_ of the original model. As mentioned above, we tested using a current that resulted in 19 spikes in 500 ms. We saw that there was an improvement in DBLO to ∼16 mV (Fig 2B).

**Fig. 2.**
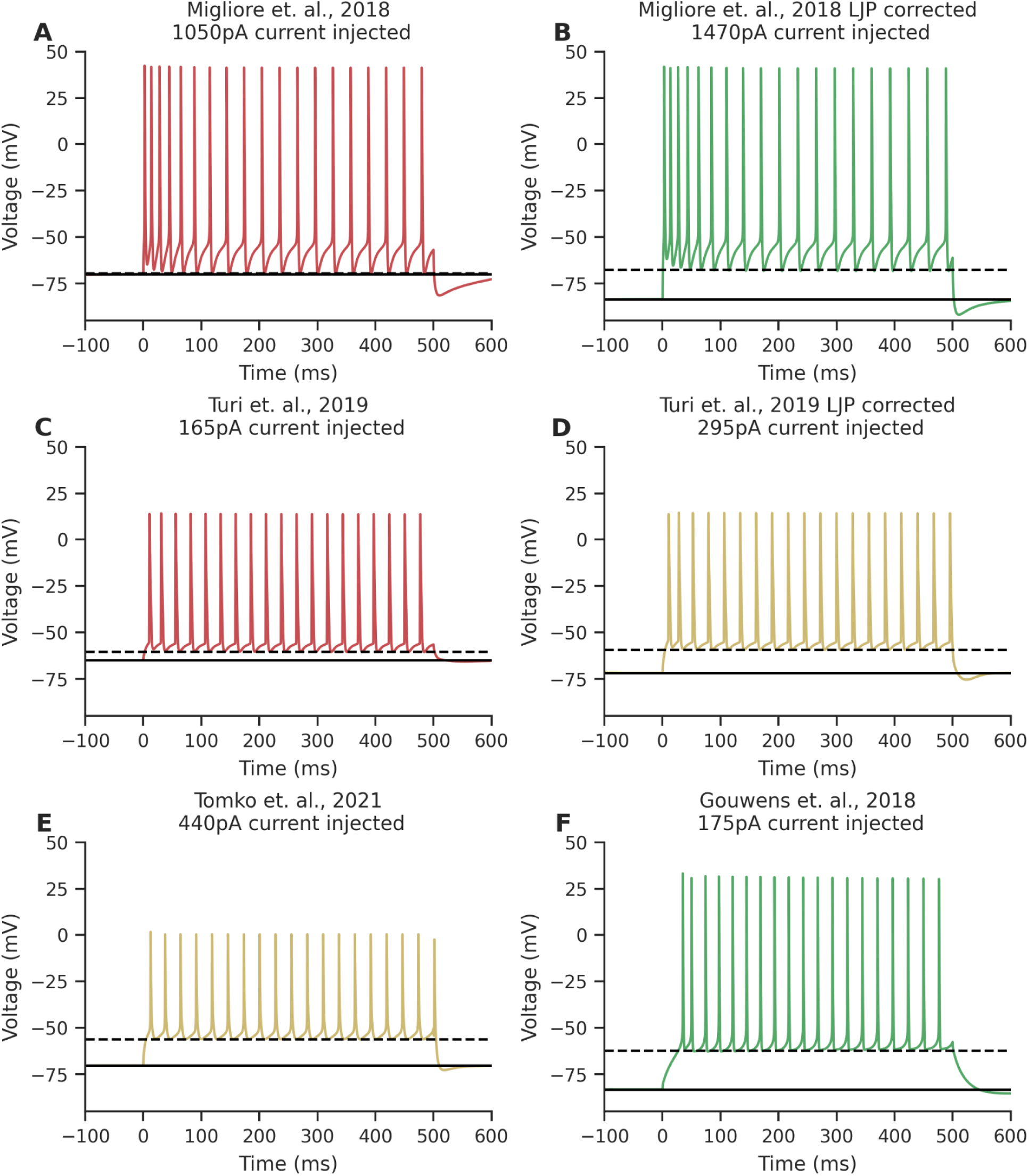
DBLO in a few state-of-the-art pyramidal neuron models. The injected current was chosen so as to get 19 spikes in 500 ms to match the representative experimental recording shown in Fig. 1A. For representational purposes, models with low DBLO of < 10 mV are colored red, models with moderate DBLO of 10 mV to 14.3 mV are colored yellow, and models with high DBLO of > 14.3 mV are colored green. **(A)** A rat CA1 pyramidal neuron model from (R. Migliore et al., 2018) showing a low DBLO of 1 mV. **(B)** A modified (R. Migliore et al., 2018) model, which accounts for LJP. Note the high DBLO of 16 mV. **(C)** Rat CA1 pyramidal neuron model from (Turi et al., 2019) with a DBLO of 4.7 mV. **(D)** A modified (Turi et al., 2019) model that accounts for LJP. Note the moderate DBLO of 13 mV. **(E)** Rat CA1 pyramidal neuron model from (Tomko et al., 2021) showing moderate DBLO of 14 mV. **(F)** Rat cortical pyramidal neuron model from (Gouwens et al., 2018) showing high DBLO of 21 mV.

Similarly, the rat CA1 pyramidal neuron model from (Turi et al., 2019), after adjusting for the number of spikes, also showed a low DBLO of 4.7 mV (Fig. 2C). This value increased to 13 mV after we compensated the model for LJP (Fig. 2D).

In contrast, the rat CA1 pyramidal neuron model from (Tomko et al., 2021) displayed a moderate DBLO of 14 mV even in the absence of LJP correction (Fig. 2E). This is one of the few conductance-based CA1 pyramidal models in our test set that exhibited DBLO within the experimental range of 11.6 mV to 27.6 mV (at 150 pA).

One of the rat cortical pyramidal neuron models from (Gouwens et al., 2018) showed a high DBLO of 21 mV when they corrected their data for LJP. Although this model has a high DBLO and other electrophysiological properties well within the experimental range (Gouwens et al., 2018), the morphology and kinetics of ion channels used to build these models were taken from cortical pyramidal neurons instead of CA1 pyramidal neurons. The DBLO in the various models in Fig. 2 is summarized in Table 1. Thus, our model survey showed that high DBLO can exist in HH-type conductance-based models, and LJP correction increases DBLO. We next examined why LJP correction is necessary for high DBLO models.

**Table 1:** Selected pyramidal neuronal models and their DBLO levels.

| Source | Original DBLO | Correction | Final DBLO |
| --- | --- | --- | --- |
| (R. Migliore et al., 2018) | 1 mV | $E_m$ reduced by 50 mV | 16 mV |
| (Turi et al., 2019) | 4.7 mV | $E_m$ reduced by 35 mV | 13 mV |
| (Tomko et al., 2021) | 14 mV | NA | NA |
| (Gouwens et al., 2018) | 21 mV | NA | NA |

### Liquid Junction Potential correction of experimental data is necessary for building models with high DBLO

We illustrate that the experimental recording needs to be corrected for LJP (sometimes referred to as pipette tip potential) before using this data for modeling. Fig. 3A demonstrates the origin of LJP. LJP arises due to ion concentration differences between the intracellular and external buffer solutions used in experiments (Neher, 1992; Zielen, 1963). The typical values of LJP can be anywhere between 0 mV and 20 mV, depending on the solutions used. Some studies correct this offset when they report membrane potentials. Other studies instead choose only to report the LJP offset values, while some studies ignore LJP altogether.

**Fig. 3.**
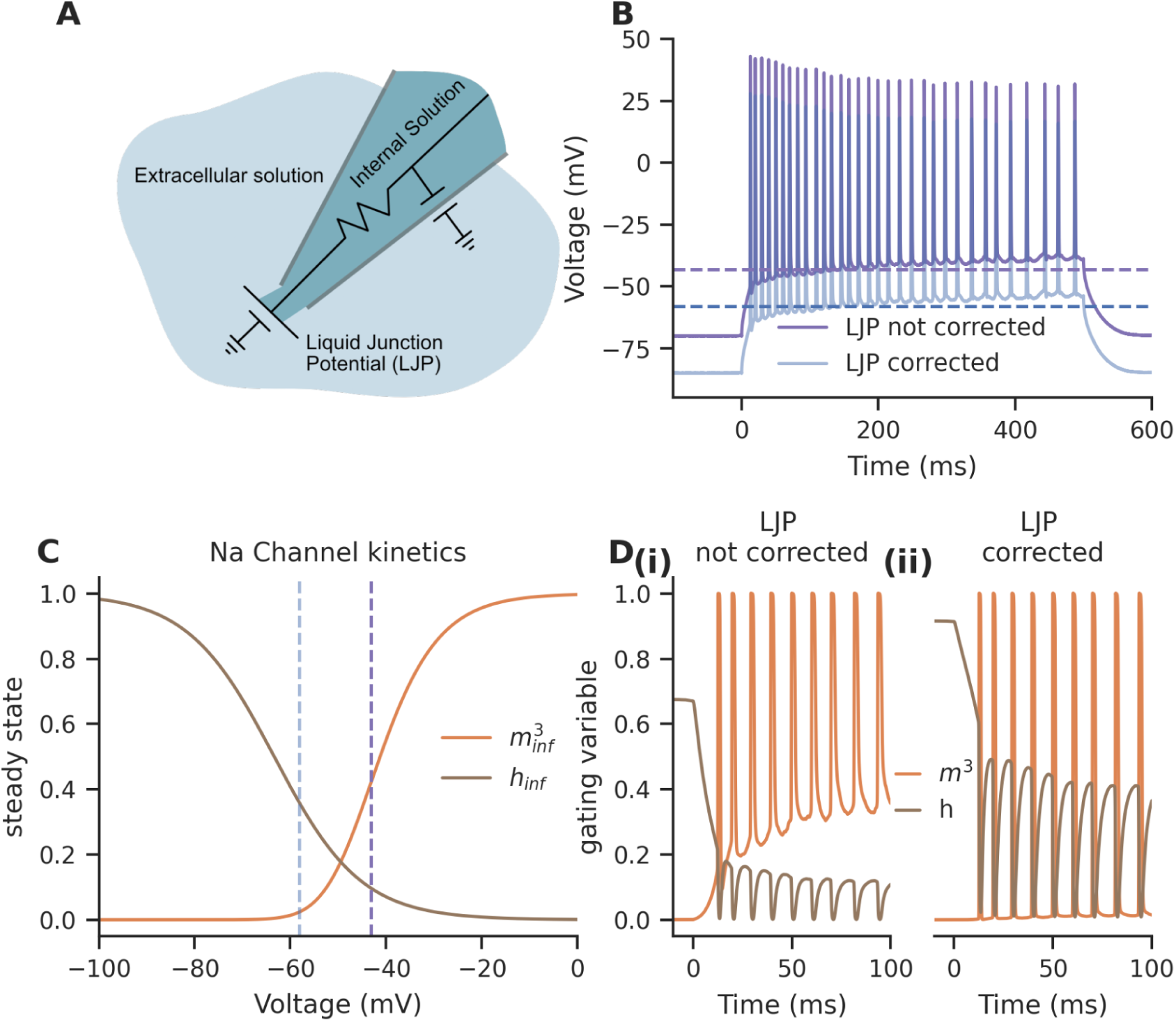
Not accounting for LJP leads to Na^+^ channel inactivation at high DBL. **(A)** Illustration showing the source of LJP. LJP emerges when liquids of different electrolytes come into contact. **(B)** A representative experimental recording at 300 pA current injection in a CA1 pyramidal neuron where LJP is accounted for (light blue) and not accounted for (dark blue). Dashed lines indicate respective DBL. **(C)** Steady-state activation and inactivation curves of Na_T kinetics. The dark blue vertical dashed line represents the DBL when 300 pA current is injected into a CA1 pyramidal neuron, and LJP is not accounted for. The light blue vertical dashed line represents the DBL when 300 pA current is injected into a CA1 pyramidal neuron, and LJP is accounted for. **(D)** The experimental recording shown in B was used to calculate the opening probabilities of the Na_T gates m and h. **(i)** When LJP was not corrected, the inactivation gate did not recover after the first AP, and further APs were not possible. **(ii)** When LJP was corrected, the inactivation gate partially recovered after the first AP, and further APs were possible.

We first calculated LJP (15 mV) from the buffers we used in the pipette and external bath (Materials and Methods) and subtracted the LJP from the raw voltage traces to get the corrected traces. Fig 3B replicates Fig 1B with and without LJP offset correction at 300 pA current injection. To understand how the Na_T channels might behave under the two scenarios, we computed the temporal evolution of its activation and inactivation variables under these two conditions. We used Na_T kinetics from (Royeck et al., 2008) (Fig. 3C) and equations 8 & 13, which describe this temporal evolution for this purpose. The inactivation gate did not recover after the first spike in the case of the trace where LJP was not corrected (Fig 3D i). However, for the LJP-corrected trace, the inactivation gate recovered to higher levels (Fig 3D ii). This was because when LJP is not corrected, the DBL is at around −40 mV (green dashed line in Fig 3B and Fig 3C). At this high potential, the steady state of the inactivation variable (h_inf_) is almost 0 (Fig 3C); thus, all the Na^+^ ion channels would be inactivated. If the LJP is corrected, the DBL is at around −60 mV, where the inactivation gate does not fully inactivate (red dashed line in Figs. 3B and 3C). Hence, without LJP correction, the models will not lead to an AP train at such a high DBL.

Having explored the role of LJP correction as a key criterion for achieving high DBLO, we next examined how a neuron’s passive and morphological properties might affect DBLO. For the remainder of the results, only LJP-corrected data is used.

### Localizing ion channels in the Axon Initial Segment results in high DBLO but low spike amplitudes

In most pyramidal neurons, the Axon Initial Segment (AIS) is the initiation center of APs and expresses major ion channels required for AP initiation (Colbert & Johnston, 1996; Kole et al., 2008). Since experimental recordings are usually taken from the soma, we hypothesized that due to the RC (resistor-capacitor) filtering property of the membrane and cytoplasmic stretch between AIS and soma, the APs generated at the AIS would experience low pass filtering. This may result in the filtering out of the trough, which is a high-frequency part of the AP shape, thus leading to increased DBLO.

To test this, we constructed two-compartment models featuring a passive soma and Na_T and K_DR channels localized in the AIS. An illustration of the morphology and the hypothesis is shown in Fig. 4A. The passive parameters, maximal conductance of Na_T (Na_T_Gbar), and maximal conductance of K_DR (K_DR_Gbar) were kept as free parameters when generating these models. The models were considered valid only if their passive properties were within the experimental range and firing frequency was within the 10 - 90^th^ percentile range of experimental data (see materials and methods for detail). We observed that, indeed, the DBLO increased with an increase in the axial resistance (RA), which is a proxy of the electrotonic distance between the AIS and soma (Fig 4B). However, apart from the DBLO increase, we also saw that the spike height decreased to non-physiological levels as RA increased (Fig. 4C). 150 pA responses of two representative models that showed high DBLO and low DBLO are demonstrated in Fig. 5D, with the high DBLO model showing extremely low and non-physiological AP amplitude.

**Fig. 4.**
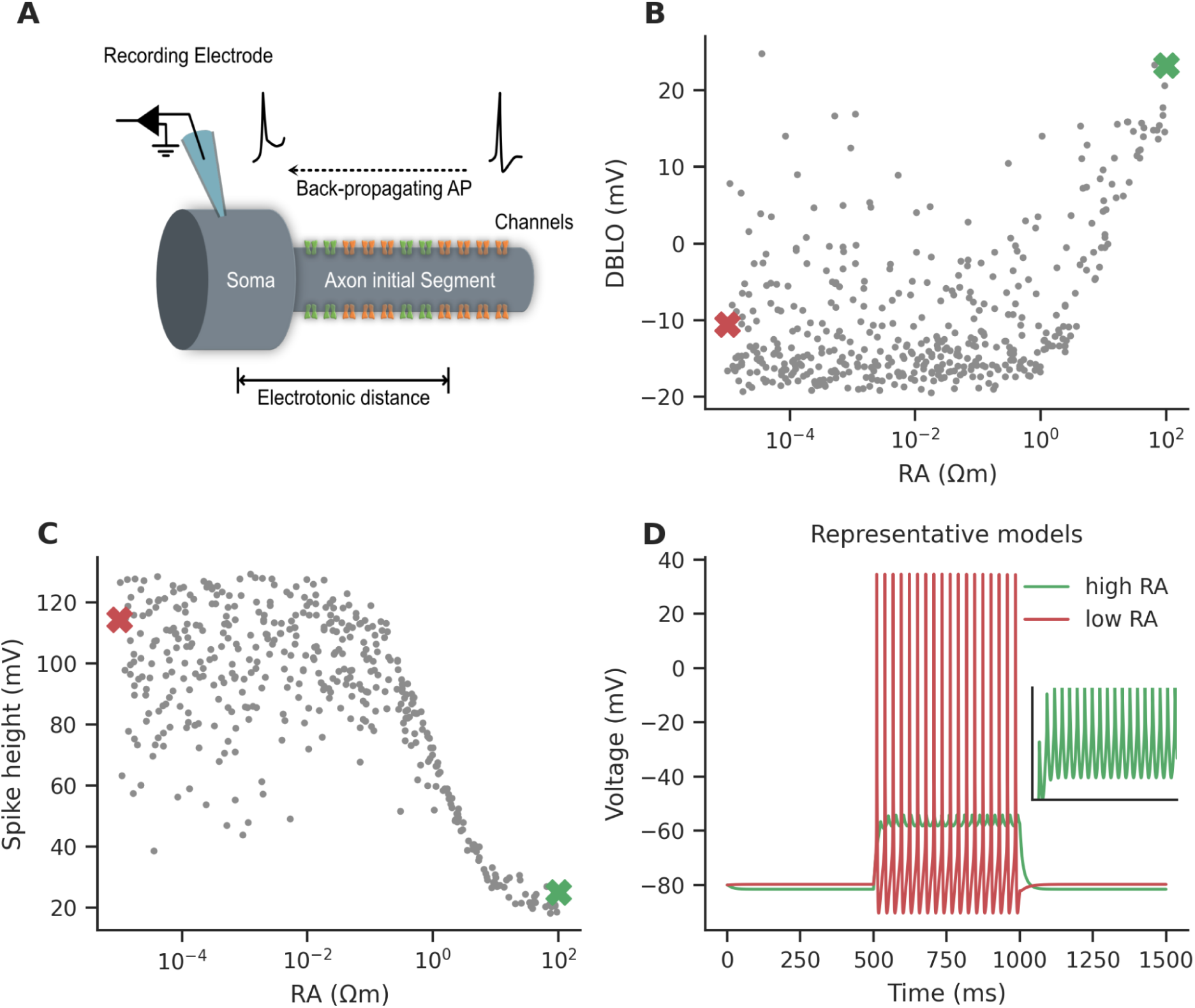
High DBLO is not due to AP initiation in AIS. **(A)** Morphology and ion channel distribution. The ion channels were inserted into the AIS section, and soma was kept completely passive. **(B)** DBLO was positively correlated with the RA (Pearson’s correlation coefficient = 0.60, p-value < 0.001). **(C)** The spike height decreased to non-physiological levels when RA was increased (Pearson’s correlation coefficient = −0.55, p-value < 0.001). **(D)** Two extreme models with the highest RA (green) and the lowest RA (red). Since the spike height of the highest RA model is too low, its APs are not visible. The APs are visible in the inset, which shows the zoomed trace. The inset features an x-axis spanning 500 ms to 1000 ms, while the y-axis extends from −59.2 mV to −54.2 mV. The highest and lowest RA models are indicated by green and red x marks, respectively, in panels B and C.

**Fig. 5.**
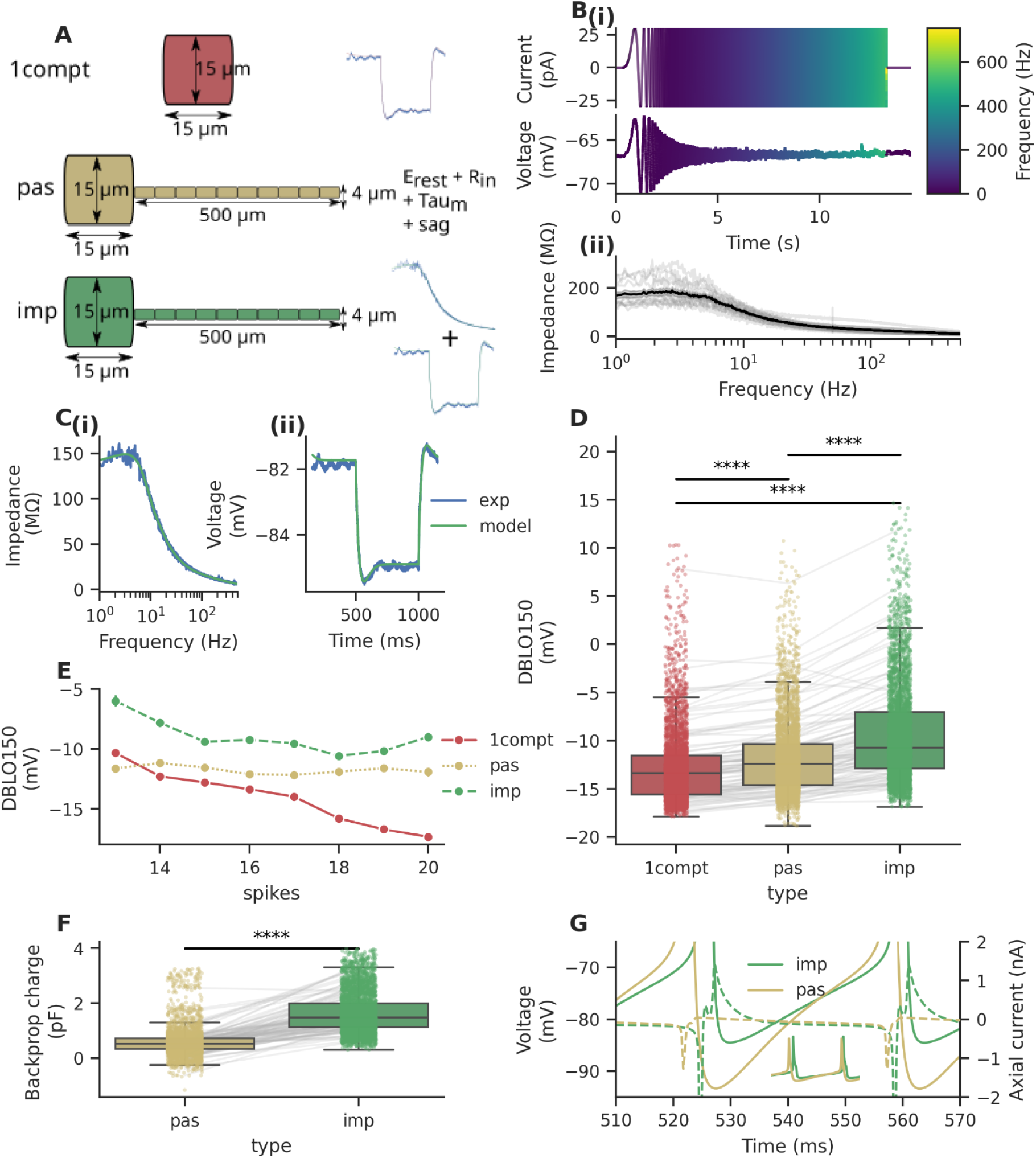
Morphology affects DBLO. **(A)** The morphologies used for generating *1compt*, *pas*, and *Imp* models, and the experimental data to which they were fitted. **(B)** Chirp response and impedance. **(i)** Injected current trace and the corresponding voltage recorded from a neuron. The color represents the sinusoidal frequency of the injected current. **(ii)** The black curve shows the mean impedance amplitude calculated from the voltage response and the current injected from all 13 recorded neurons. Gray lines represent the impedance amplitude profiles of individual neurons. **(C)** Passive fits for the representative *imp* model and its source neuron. Left: Impedance amplitude profile. Right: −25 pA response. **(D)**: DBLO comparison for all sampled models. The *imp* models had higher average DBLO than the *pas* and *1compt* models (see main text for details and statistics). **(E)** Mean DBLO comparison for the population of models separated by the number of spikes when 150pA current is injected for 500 ms. As before, *imp* models have a higher average DBLO. **(F)** The total backpropagating charge coming into the soma from the adjacent compartment between the first AP’s peak time and its trough time. *Imp* models had significantly higher charges coming into the soma during the repolarization phase (see main text for stats). **(G)** The axial current from the soma into the adjacent compartment in a pair of *imp* and *pas* models. Solid lines are the AP waveform zoomed in to show the DBLO difference between the two models. Dashed lines show the axial currents. The inset shows the zoomed out AP waveform. The peak axial currents in *imp* models are much larger than in *pas* models.

Although a physiologically faithful configuration would require an active soma, we here checked for the extreme case where the somatic channels do not influence APs. Thus, the initiation of APs in AIS alone cannot reconcile high DBLO with observed spike amplitudes. Therefore, we did not separate AIS and soma in the subsequent results and analysis. Next, we looked at whether dendritic morphology can affect DBLO.

### Multicompartment models with physiological frequency responses produce higher DBLO than single-compartment models

(Hendrickson et al., 2011) showed that the afterhyperpolarization (AHP) was shallower in models with higher dendritic resistance. Higher dendritic resistance meant the dendritic compartments charged to a higher level during AP depolarization, and subsequently, during somatic repolarization, dendritic axial currents fed into the soma, causing shallower AHP. Thus, we hypothesized that a similar mechanism may affect DBLO. To explore this, we systematically tested groups of models with varying levels of electrotonic coupling between the soma and dendrites, ranging from completely isolated somas to experimentally calibrated coupling.

We first developed three families of subthreshold models: (1) single-compartment models (*1comp*) that match the passive features of the recorded CA1 pyramidal neurons, (2) multi-compartment ball-and-stick models (*pas*) that also match these passive features, and (3) multi-compartment ball-and-stick models (*imp*) that match both the passive features and the subthreshold impedance amplitude of the recorded CA1 pyramidal neurons (Fig. 5A). All models included the HCN channel to account for the voltage sag in recordings (Albertson et al., 2017; Yi et al., 2016). For passive features, we captured the following experimental passive properties - E_rest_, R_in_, C_in_, charging time constant, and sag. For *pas* models, these properties were directly fitted, while for 1comp and *imp* models, they were indirectly captured by fitting to a hyperpolarizing −25 pA current pulse (Fig. 5A). For impedance amplitude, we injected a chirp current with a subthreshold amplitude of 30 pA, which spanned 0 Hz to 500 Hz in 13 s (material and methods). We recorded the voltage response and calculated the subthreshold impedance amplitudes from these recordings (Fig. 5B). To build *imp* models, we fitted the impedance amplitude profile of the ball-and-stick model to that of the experimentally recorded neurons. This procedure captured the electrotonic coupling between the soma and dendrites, and the passive contributions of neuronal morphology to somatic potential at various frequencies which may get ignored otherwise. Ultimately, we obtained a single *1compt* model, a single *imp* model, and ten *pas* models for each of the 13 neurons.

We then proceeded to generate active models capable of producing APs. We made model triplets for each neuron by picking the corresponding *1compt* model, *imp* model, and one of the ten *pas* models. We then incorporated Na_T and K_DR into these models such that within each triplet, the maximal conductance (Gbar) of these two ion channels and the HCN ion channel were the same. We considered the triplet valid if all three models in the triplet passed our validity criterion (materials and methods). A total of N=4174 valid active models per group were built in this manner.

We observed that *imp* models exhibited the highest DBLO, followed by *pas* models, with *1compt* models showing the lowest DBLO (Fig. 5D; mean DBLO in *1compt* = −12.75 mV, *pas* = −11.74 mV, *imp* = −9.31 mV; repeated measures ANOVA F statistic = 40.15, p-value < 0.001; paired t-test *1compt*-vs-*imp* p-value < 0.001, *1compt*-vs-*pas* p-value < 0.001, *pas*-vs-*imp* p-value < 0.001).

Very few of the *1compt* and pas models (four *1compt* and one *pas*) in our parameter sweep exhibited even moderate DBLO. *Imp* models, on the other hand, yielded 19 models with moderate DBLO. One of the *imp* models had a high DBLO of 14.64 mV. We compared the models within triplets and found that, on average, the *imp* models had 3.44 mV higher DBLO than the *1compt* model, while the biggest difference was 9.17 mV.

To remove the effects of different firing rates in the models, we also plotted the DBLO as a function of the number of spikes in response to 150 pA within 500 ms (Fig. 5E). Here, too, the *imp* models showed higher DBLO than *pas* or *1compt* models.

To delineate the mechanism behind this increase in DBLO in *imp* models, we calculated for each model the total charge that enters the soma during the repolarization phase of an AP induced by 150 pA current injection. We saw that *imp* models had significantly higher amounts of this total backpropagating charge (Fig. 5F; 1.64 ± 0.74 pF for *imp* models and 0.58 ± 0.50 pF for *pas* models; paired t-test p-value < 0.001). This charge depolarizes the soma and leads to higher DBLO. To illustrate it further, we plotted axial currents into the soma during an AP in a representative pair of *imp* and *pas* models. The two models had the same Gbars and passive properties as described earlier. The axial currents during the repolarization phase can be seen to be at much higher levels in the *imp* models than in the *pas* models (Fig. 5G, dashed traces). These calculations showed that morphology contributes significantly to the DBLO. Next, we investigated the role of ion channel kinetics in this process.

### High K_DR reversal potential does not explain high DBLO

Next, we wanted to see if K^+^ channels have any effect on DBLO. We first checked if the reversal potential of K_DR had any impact on DBLO. We hypothesized that a more depolarized reversal potential (E_rev_) would result in a low driving force of the K_DR channel and, hence, would lead to high DBLO. To check this, we did a stochastic parameter search with Na_T_Gbar, K_DR_Gbar, and K_DR E_rev_ as free parameters. We indeed found that high K_DR E_rev_ resulted in high DBLO (Fig. 6A). However, high DBLO at 150 pA was achieved only when the K_DR E_rev_ was at non-physiological levels of above −70 mV. Further, we expect that DBLO should increase with higher current, as shown above (Fig. 1C) and in an experimental representative neuron recording at 150 pA and 300 pA (Fig. 6A). However, the models with high K_DR reversal potentials did not show this current-dependent increase in DBLO. The model with the lowest DBLO is demonstrated in Fig. 6B, and the model with the highest DBLO is shown in Fig. 6C. These two models show minimal increase in DBLO when injected current is increased, which is inconsistent with the recordings.

**Fig. 6.**
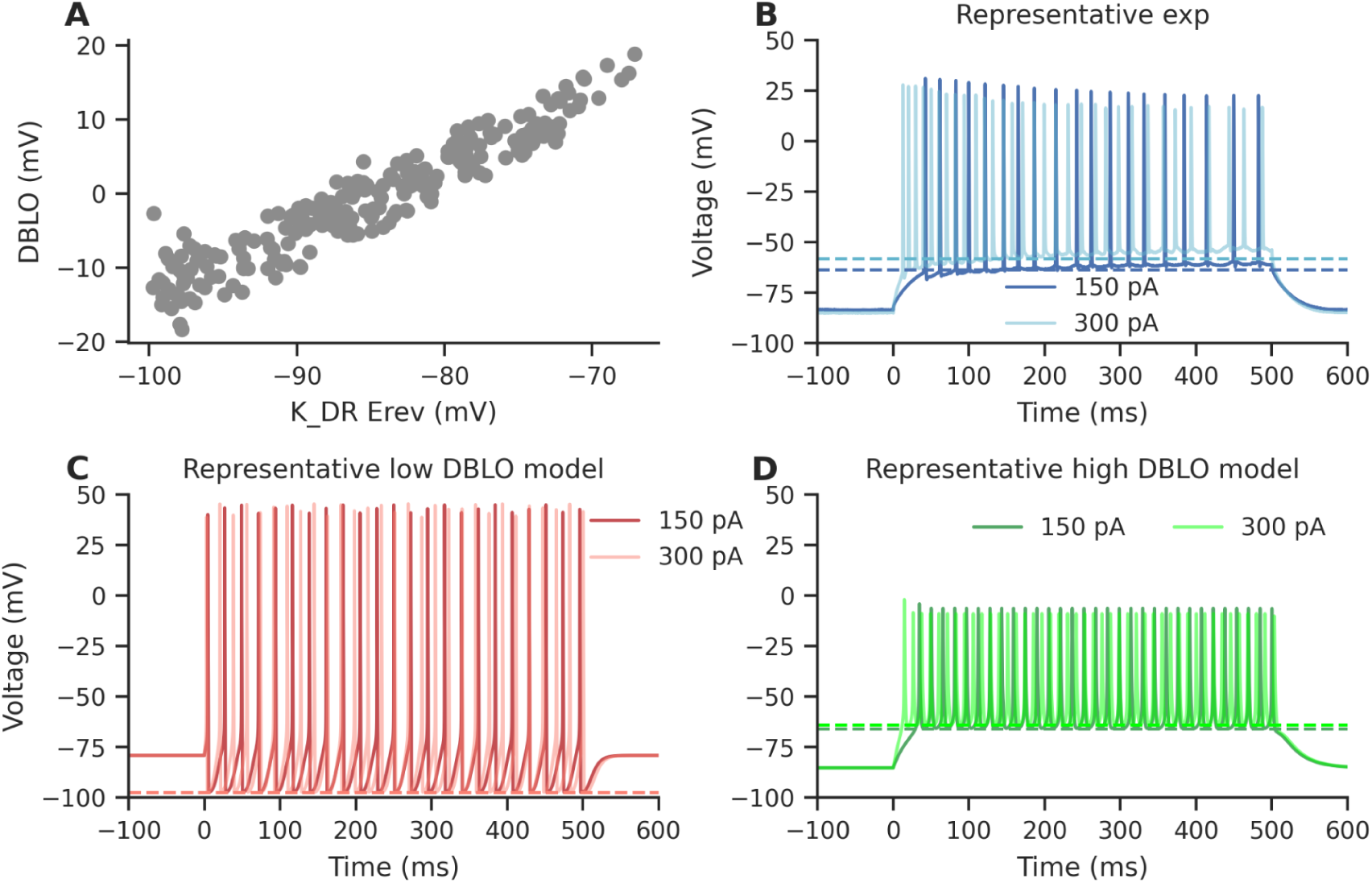
Unphysiologically high E_rev_ of the K^+^ channel gives rise to high DBLO. **(A)** DBLO increases with higher E_rev_ for K_DR but requires unphysiologically high E_rev_ to obtain high DBLO (Pearson’s correlation coefficient = 0.94, p-value < 0.001). **(B)** A representative experimental recording demonstrating that DBLO increases with increasing injection current. **(C)** A low DBLO model that does not show a current-dependent increase in DBLO. **(D)** A high DBLO model with shifted E_rev_ for K_DR that does not show a current-dependent increase in DBLO.

### Fast potassium channel deactivation kinetics result in higher DBLO than slow kinetics

We hypothesized that if the deactivation of K_DR is fast, K_DR would quickly close during the repolarization phase of an AP. This would result in faster AP termination and, hence a higher DBLO. Thus, to test this hypothesis and to explore the role of potassium channel kinetics in the high DBLO phenomenon, we constructed a thousand single-compartment models that had their passive features and firing frequency within the experimental range (Materials and Methods). We first took the thirteen *1compt* models constructed earlier that matched the experimental −25 pA voltage response. Model morphology and the equivalent electrical circuit are given in Fig. 7A. Using these thirteen models as a base, we built 677 valid active models. In constructing these models, Na_T_Gbar, K_DR_Gbar, and K_DR time constants were kept as free parameters (see eqn 22-24 for details). Figure 7B illustrates the parameters that define the tau curve of the K_DR channel.

**Fig. 7.**
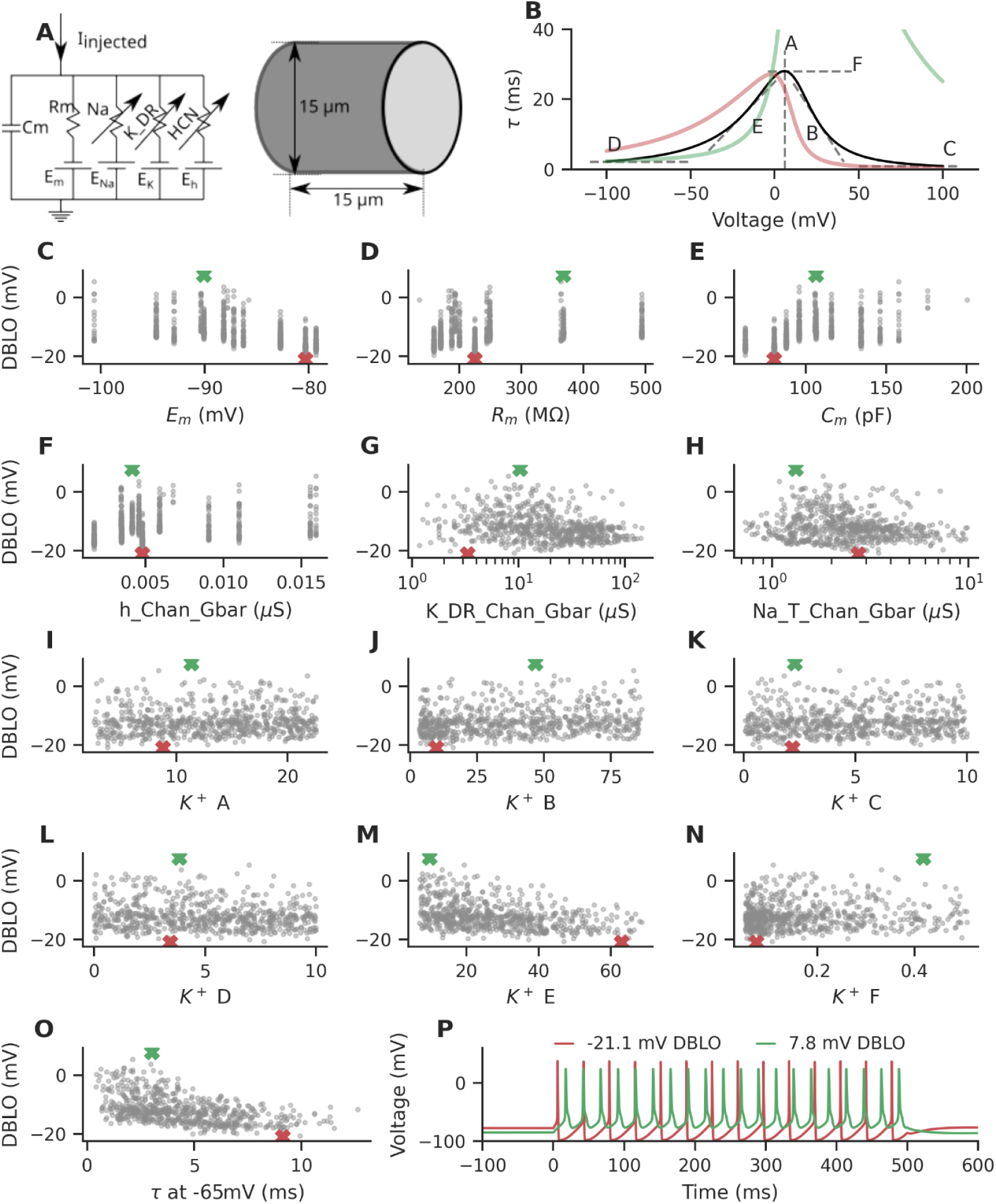
K_DR deactivation at hyperpolarized membrane potentials contributes significantly to DBLO. **(A)** Equivalent circuit diagram of the model. A single-compartment model with 15 µm length and 15 µm diameter was used. **(B)** The parameterized K_DR kinetics. Parameter A controls the peak location, parameter B controls the slope of the curve at high potentials, parameter C controls the minimum time constant at high potentials, parameter D controls the minimum time constant at low potentials, parameter E controls the slope of the curve at low potentials, and parameter F controls the peak height. Green and red curves are the K_DR kinetics of models presented in panel P. The green curve has been truncated at the top to allow better visualization. **(C-N)** The DBLO in models plotted against parameter values. Green and red X marks are the parameters associated with models presented in panel P. **(O)** DBLO vs. time-constant tau at −65 mV of K_DR deactivation. A moderate negative correlation is seen between tau and DBLO, details in text and Table 2. Green and red X marks represent the tau at −65 mV of K_DR kinetics associated with models presented in panel P. **(P)** Response to 150 pA of the model that expresses the lowest DBLO (colored in red) and of the model that expresses the highest DBLO (colored in green). The parameters corresponding to these models are shown as red and green X marks, respectively, in panels B-O.

**Table 2:** Pearson’s correlation coefficients for the parameters in models used to explore K_DR kinetics associated with high DBLO. The significance level was chosen to be 0.0056 after Bonferroni correction.

| Parameter | Pearson's correlation coefficient | p-value |
| --- | --- | --- |
| $E_m$ | -0.54 | < 0.001 |
| $R_m$ | 0.26 | < 0.001 |
| $C_m$ | 0.46 | < 0.001 |
| $h_{Chan\_Gbar}$ | 0.23 | < 0.001 |
| $K_{DR\_Chan\_Gbar}$ | -0.26 | < 0.001 |
| $Na_T\_Chan\_Gbar$ | -0.17 | < 0.001 |
| A | 0.05 | 0.172 |
| B | 0.17 | < 0.001 |
| C | 0.05 | 0.239 |
| D | -0.14 | < 0.001 |
| E | -0.37 | < 0.001 |
| F | 0.07 | 0.07 |
| K_DR tau at -65 mV | -0.48 | < 0.001 |

The passive parameters - Em, Rm, and Cm - showed moderate negative, weak positive, and moderate positive correlation with DBLO respectively (Fig. 7C-E). The maximal conductance of HCN, K_DR, and Na_T channels showed weak positive, weak negative, and very weak negative correlation (Fig. 7F-H). Among the free parameters associated with K_DR kinetics (Figs. 7I-N), we observed a significant but weak negative correlation between E and DBLO (Fig. 7M; Table 2). For all other parameters, we either observed very weak or no significant correlation (Table 2). Low values of E correspond to low tau (fast deactivation kinetics) at low voltages, as evident in Fig. 7O, where a moderate negative correlation between DBLO and tau at −65 mV in models can be seen. To illustrate the point, we compare 150 pA current injection traces from two representative models in Fig. 7P. The model shown in green has a DBLO of 7.8 mV, while the model shown in red has a DBLO of −21.1 mV. The parameters associated with these models are marked in green and red, respectively, in Figs. 7B-O. Notably, the tau at −65 mV of the model with lower DBLO is indeed higher than that of the model with higher DBLO, reinforcing our hypothesis (Fig. 7O).

The Pearson’s correlation coefficient for the free parameters vs. DBLO are listed in Table 2.

### Na_T kinetics has a significant impact on DBLO

To check whether Na_T kinetics might have any effect on DBLO, we made three groups of active models, which we refer to as *Gou*, *Roy*, and *Mig* models. In all three groups of models, we used the subthreshold *1compt* models generated previously, and the default slow K_DR kinetics from (R. Migliore et al., 2018).

For *Gou* models, we incorporated Na_T kinetics from (Gouwens et al., 2018) which are based on (Hay et al., 2011), which in turn are derived from the experimental kinetics from neocortical layer 5 pyramidal neurons of rats (Colbert & Pan, 2002). For *Roy* models, the Na_T steady-state curve was taken from experimental data from mouse CA1 pyramidal neurons (Royeck et al., 2008). Since their study did not provide tau curves, we adopted the tau curves from (Hay et al., 2011). For the *Mig* models, we implemented Na_T kinetics from (R. Migliore et al., 2018), which were in turn adapted from (M. Migliore et al., 1999) with slight modifications to the experimental kinetics originally recorded from rat CA1 pyramidal neuron dendrites (Magee & Johnston, 1995). The kinetics are shown in Fig. 8A. For each model group, we varied the Gbar of the Na_T and K_DR channels while picking one of the thirteen *1compt* models at random. We tested 6500 model configurations per group and rejected models that did not have active properties within the experimental range (see materials and methods).

**Fig. 8.**
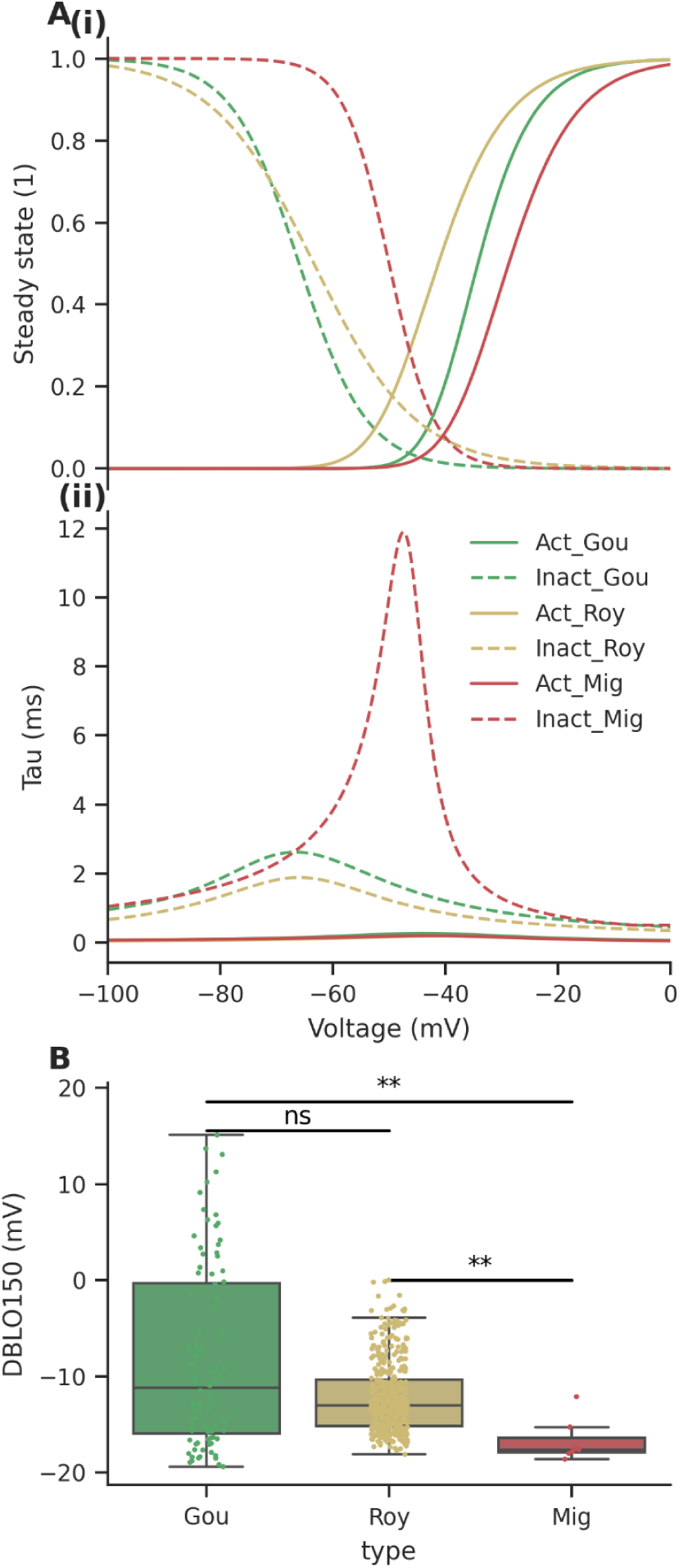
Effect of Na_T kinetics on DBLO. **(A)** Na_T kinetics from three different sources: (Gouwens et al., 2018; R. Migliore et al., 2018; Royeck et al., 2008). **(i)** The steady state curves of the activation (solid lines) and inactivation (dashed lines) gates. **(ii)** The tau curves. **(B)** DBLO was significantly different between models made using the three Na_T kinetics (Kruskal-Wallis H-test, statistic = 13.4, p-value = 0.0012). Na_T kinetics from *Gou* showed highest mean DBLO of −8.3 ± 9.4 mV, followed by *Roy*, with mean DBLO of −12.2 ± 3.7 mV, and *Mig*, with mean DBLO of −16.7 mV ± 2.3 mV. The differences in medians between the *Gou* and *Roy* models were not significant (Dunn’s post-hoc test p-value = 0.3), while the *Gou* vs. *Mig* (Dunn’s post-hoc test p-value = 0.001), and *Mig* vs. *Roy* (Dunn’s post-hoc test p-value = 0.0043) were significant after Bonferroni correction.

We saw that there was a significant difference in the median DBLO levels of the three groups of models (Fig. 8B). The *Gou* models had the largest mean DBLO. It was also the only group with a few (5) moderate DBLO models out of a total of 406 valid models (one model with high DBLO). The post-hoc Dunn test revealed that there was no significant difference between the *Gou* and *Roy* groups. At the same time, the DBLOs were significantly different in the pairwise comparisons between *Gou* vs. *Mig*, and *Mig* vs. *Roy* (statistical data in Fig. 8 legend).

Up until now, we have seen that independently optimizing morphology, K_DR, and Na_T kinetics could significantly increase the DBLO compared to using default values. Next, we show that combining these factors gives rise to even higher DBLO models. We also explore the implications of considering DBLO while building conductance-based models.

### Multiple factors together contribute to high DBLO

We showed above that multiple factors contribute to DBLO, mainly LJP correction, the neuron’s subthreshold impedance profile, K_DR deactivation kinetics, and Na_T kinetics. Although these factors had a significant effect on DBLO, they could not separately account for the high DBLO seen in our recordings. In particular, only one model with a physiological subthreshold impedance profile and one model with (Gouwens et al., 2018) Na_T kinetics approached the observed experimental DBLO levels. A summary of these findings is presented in Table 3.

**Table 3:** Summary of various potential factors that were tested.

| Factors | Contribution | Maximum DBLO in simulations | Comments |
| --- | --- | --- | --- |
| LJP correction | High | NA | Na channels inactivate in the absence of LJP correction |
| Contribution from AIS | Low | 24.83 mV | Non-physiologically low spike amplitudes accompany high DBLO |
| Dendritic morphology or the Impedance amplitude profile | High | 14.65 mV | Models that matched experimental impedance amplitude profiles had higher DBLO |
| K_DR $E_{rev}$ | Low | 18.87 mV | Non-physiological $E_{rev}$ needed to achieve high DBLO |
| K_DR kinetics | High | 7.8 mV | The deactivation time constant of the primary potassium ion channel is negatively correlated with DBLO |
| Na_T kinetics | High | 15.12 mV | Incorporating Na_T kinetics from (Gouwens et al., 2018) gave high DBLO |
| Compound effects of all the factors | High | 18.1 mV | The combined factors give a high DBLO close to the experimental mean |

To assess whether combining these factors would yield high DBLO models, we combined these factors into a single group of models. We used the following configuration:

1. We used the previously generated subthreshold *imp* models as a base.
2. We incorporated a fast-deactivating K_DR channel with the kinetics E parameter reduced by half compared to the K_DR kinetics from (R. Migliore et al., 2018). To compensate for the unintended changes to the time-constant curve, F was multiplied by 1.1 and A shifted right by 0.006. This reduced the tau at −65 mV from 4.5 ms to 1.5 ms making deactivation at −65 mV ∼3 times faster.
3. We used an elevated K_DR E_rev_ of −90 mV instead of −100 mV.
4. We adopted Na_T kinetics from (Gouwens et al., 2018) instead of from (Royeck et al., 2008)

A stochastic parameter search on Gbars of the Na_T and K_DR channels and these modifications gave us 1171 valid models out of the 65000 combinations tested (see materials and methods). Henceforth, we will refer to these 1171 valid models as valid *unified* models. Among these, 46 models exhibited high DBLO of more than 14.3 mV. Remarkably, one of the models had a high DBLO of 18.1 mV, which was close to the experimental mean of 18.8 mV. The 150 pA response of this model is shown in Fig. 9A.

**Fig. 9.**
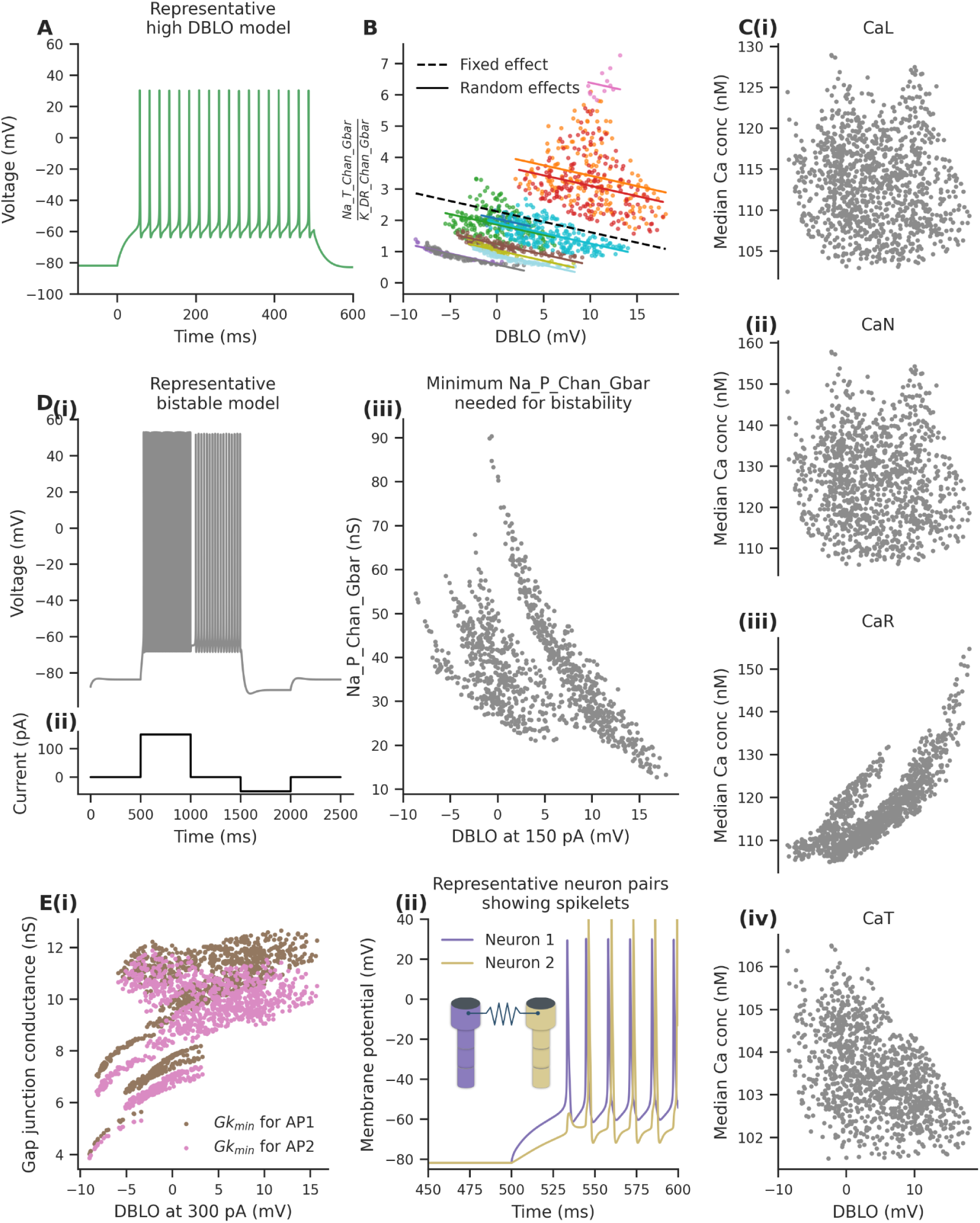
Not considering high DBLO in models may lead to incorrect quantitative and qualitative predictions. **(A)** Representative high DBLO model. The model used a ball and stick morphology that matched the subthreshold impedance amplitude profile of recorded neurons, had fast K_DR kinetics, and used Na kinetics from Gouwens et al., 2018. **(B)** The ratio of maximal conductance of Na_T and K_DR is negatively correlated with DBLO levels. Models with different base subthreshold *imp* models show different intercepts (shown here with separate colored dots). Note that only 11 groups are present, as two of the base subthreshold *imp* models did not give rise to any valid active models. **(C)** Different calcium conductance types were incorporated into valid unified models. The median calcium concentration during a train of APs induced by 300 pA current injection was measured in these models. The median internal Ca concentration when either CaL or CaN were present in the model did not show any correlation. The similarity in the CaL and CaN plots arises from their resulting median internal calcium concentrations being equivalent, despite the distinct underlying dynamics of each. CaR and CaT led to a positive and negative correlation, respectively. **(D)** Na_P channel imparts bistability to valid *unified* models. **(i)** A representative bistable model. Note that the model fires even in the absence of current injection between 1500 - 2000 ms. **(ii)** The current trace injected to check for bistability. **(iii)** Minimum Na_P maximal conductance needed for valid *unified* models to fire in a bistable mode correlated negatively with DBLO in these models (Pearson’s correlation = −0.53). **(E) (i)** The minimum gap junction conductance (Gk) needed for the induction of 1st and 2nd AP in a post-synaptic neuron is positively correlated with DBLO. **(ii)** At Gk less than this minimal conductance, an AP in the presynaptic neuron (Neuron 1) induces a spikelet in the post-synaptic neuron (neuron 2), as can be seen in a representative simulation with a gap junction Gk of 10.9 nS. The inset provides an illustration of gap junction connection between neuron 1 and neuron 2.

### Models with DBLO have constrained conductance ranges

Having assembled a library of high DBLO models, we next asked if they exhibited functionally distinct and computationally relevant properties compared to low DBLO models. We first asked if channel distributions were distinct.

To explore whether Na_T_Gbar and K_DR_Gbar show any dependence on DBLO levels, we plotted the ratio between their maximal conductances against DBLO in the 1171 *unified* models we built in the preceding section. We designed a linear mixed-effect model (LMM) to check for correlations between DBLO and the ratio of Na_T_Gbar and K_DR_Gbar. The LMM included DBLO as a fixed effect and accounted for random effects across the 13 base subthreshold *imp* models. We observed a significant negative association between DBLO and Gbar_ratio (β = −0.067, p-value < 0.001), suggesting that higher DBLO values correspond to a decrease in the Gbar_ratio (Fig 9B). This implies that models built without considering DBLO may give different quantitative predictions about the maximal conductance levels of ion channels as compared to models that were constructed while keeping DBLO as a feature. This correlation may be biologically significant as the Gbar_ratio more than quadrupled in models with DBLO of 15 mV compared to models with DBLO of −10 mV (Fig. 9B).

### DBLO levels are correlated with internal calcium concentration in models

We hypothesized that DBLO might affect the internal calcium concentration. We used our set of unified 1171 valid models and incorporated different calcium conductance types in these models. The kinetics and maximal conductance of these conductances were taken from previous modeling studies (R. Migliore et al., 2018; Upchurch et al., 2022). We measured the internal calcium concentration in these models and checked for any correlations with DBLO. We found no correlation when either L-type (Pearson’s correlation coefficient = −0.075; p-value = 0.010; Fig. 9C i) or N-type (Pearson’s correlation coefficient = −0.070; p-value = 0.017; Fig. 9C ii) calcium channels were incorporated. When we incorporated R-type and T-type calcium conductances, we found positive (Pearson’s correlation coeff 0.80; p-value < 0.001; Fig. 9C iii) and negative (Pearson’s correlation coeff −0.47; p-value < 0.001; Fig. 9C iv) correlations between internal calcium concentration and DBLO respectively. This implies that low DBLO models might incorrectly estimate levels of internal calcium levels or might need non-physiological levels of maximal calcium channel conductances to compensate. This, in turn, might affect many calcium-based plasticity mechanisms (Chindemi et al., 2022; Graupner & Brunel, 2012; Zucker, 1999).

### Bistable firing occurs in high DBLO models at lower levels of Na_P

We next asked if there were firing pattern inconsistencies that might arise from not having DBLO as a feature while building models. To do this, we looked at firing bistability. We hypothesized that certain ion channels might need to be included at very high non-physiological levels in low DBLO models to observe their physiological effects. To confirm this, we added Na_P channels in our valid *unified* models. Using a binary search method, we found the minimum Na_P_Chan_Gbar needed for a model to exhibit bistability (see materials and methods). A representative bistable model is shown in Fig. 9D i. Firing bistability was defined as the ability of a neuron to switch between tonic firing and silent mode when transient depolarizing and hyperpolarizing currents are injected, respectively. The current trace injected for this purpose is shown in Fig. 9D ii.

We found that the minimum Na_P_Chan_Gbar needed to turn a model bistable was negatively correlated with DBLO (Fig. 9D iii; Pearson’s correlation coefficient = −0.53). The high DBLO models (DBLO > 14.3 mV) significantly lower Na_P_Chan_Gbar to turn bistable than low DBLO models (DBLO < 10 mV) (High DBLO models − 17.7 ± 2.7 nS; Low DBLO models - 39.9 ± 10.7 nS; Welch’s t-test p-value < 0.001). In comparison, studies have found the somatic Na_P_Gbar in non-bistable neurons to be between 1 - 10 nS (Shvartsman et al., 2021; Yue et al., 2005). This implies that if one does not consider DBLO while building models, they might need a higher amount of Na_P_Chan_Gbar to see firing bistability, which might result in unintended effects on other Na_P mediated properties of the models such as subthreshold synaptic amplification (Hsu et al., 2018), and firing threshold (Deng & Klyachko, 2016).

### The minimal gap junction conductance needed for AP transfer from one neuron to another is correlated with DBLO

As a final examination of the relevance of appropriate DBLO on circuit function, we asked if it might have any effects on gap-junction coupled firing. Axo-axonic gap junctions between CA1 pyramidal neurons have been found to be involved in the generation of very fast oscillations (Schmitz et al., 2001; Traub et al., 1999). We hypothesized that DBLO levels might be correlated with the minimal conductance of these gap junctions needed for an AP in one neuron to induce an AP in another neuron connected via a gap junction. To test this, we used the 1171 valid unified models and connected the soma of two instances of each model via bidirectional gap junctions. Since adding a gap junction decreased the input resistance of the neuron, 150 pA did not induce any spiking. Thus, we used 300 pA to induce spiking in the soma of one instance and recorded it from the soma of the post-synaptic neuron. For each model, using binary search, we calculated the minimum gap junction conductance needed for the 1st and 2nd spikes in the AP train to be induced in the soma of the post-synaptic neuron. We found that this minimum gap junction conductance was positively correlated with DBLO for both the 1st (Pearson’s correlation coefficient 0.67; p-value < 0.001) and 2nd spike (Pearson’s correlation coefficient 0.58; p-value < 0.001) (Fig. 9E i). A possible explanation for this positive correlation is that models with high DBLO also exhibit a higher AP voltage threshold (Supplemental Fig. S1A). This is likely due to a lower ratio of Na_T_Gbar to K_DR_Gbar in high DBLO models as shown in Fig. 9B. With a higher AP threshold, a greater gap junction conductance is required to reach the necessary depolarization for AP initiation.

Interestingly, we found that the minimum gap junction conductance needed for the induction of the 2^nd^ spike was lower than that for the 1^st^ spike. This led to the induction of spikelets at certain gap junction conductances (1^st^ AP in Fig. 9E ii). Studies have proposed axo-axonic gap junctions to be one of the sources of these spikelets (Michalikova et al., 2020). One caveat is that, since our models did not have axons, we directly connected the somatic compartments. Future studies that use more morphologically accurate models of the axo-axonic gap junction might yield more accurate quantitative predictions regarding the gap junction conductance necessary for the spread of AP through gap junctions.

Thus, not considering DBLO may give incorrect predictions about the gap junction maximal conductance that allows spikelets to occur. This, in turn, could play a role in our understanding of gap junction-mediated high-frequency oscillatory events (Traub et al., 1999; Traub & Bibbig, 2000).

## Discussion

In this study, we have used models and experiments to identify mechanisms that underlie a prominent feature of neuronal firing, the DBLO. Our findings indicate that four factors substantially contribute to DBLO: LJP correction, the subthreshold impedance amplitude profile of models, fast deactivation kinetics of K_DR, and Na_T kinetics. We show that the omission of DBLO as a constraint while building models may lead to incorrect estimates of levels of ion channels as well as incorrect predictions about the role of ion channels in several cellular functions. Such quantitative details can be biologically and medically important, such as during seizures (Blumenfeld et al., 2009; Quraishi et al., 2019; Speca et al., 2014). Our analysis revealed that while none of the factors alone were sufficient to generate large numbers of high DBLO models, their combined effects successfully produced many models with DBLO values within the experimental range. This suggests that the high DBLO observed in experimental recordings likely arises from a combination of these factors.

In our minimal models, we focused on a limited set of ion channels—specifically Na_T, K_DR, and HCN—to identify key contributors to DBLO levels. However, incorporating additional physiological channels, such as A-type and SK-type potassium channels, could indirectly support action potential trains at even higher DBLO by enhancing Na_T recovery from inactivation through increased inter-spike intervals, thereby sustaining stable firing at elevated DBLO levels. The A-type potassium channels, known for their fast deactivation kinetics and significant contribution to potassium currents in CA1 pyramidal neurons (Martina et al., 1998), are a strong candidate for the fast-deactivating potassium channel underlying the high DBLO phenomenon observed in our simulations.

### Tighter constraints for morphological simplification and passive properties

Subthreshold impedance amplitude profiles have been used previously to study the resonance property of neurons (Binini et al., 2021; Dhupia et al., 2015; Moradi Chameh et al., 2021; Narayanan & Johnston, 2007). Since a neuron’s subthreshold resonance frequency is between 0-10 Hz (Hutcheon et al., 1996; Matsumoto-Makidono et al., 2016; Ulrich, 2002), these studies have used chirp stimuli that span 0 Hz to ∼50 Hz. To the best of our knowledge, ours is the first study to use chirp stimuli (Fig. 5) that span 0 Hz to 500 Hz to measure subthreshold impedance amplitude profiles and to use this to constrain multicompartment models. (Rall, 1969) has previously shown that multicompartment neurons exhibit several time constants. The faster of these time constants affect the rapid depolarization and repolarization of soma during an AP as they are of the same timescale as an AP. Thus, the morphology of a neuron indirectly affects the AP shape and DBLO of a neuron, which we have captured by measuring the subthreshold impedance amplitude.

We find that this constraint over the subthreshold impedance amplitude works well even in our simplified models in which the dendritic section is a composite of all the neurites - proximal, apical, distal, as well as axon. The caveat is that this simplification gives much lower electrotonic lengths (0.45 ± 0.18) of the subthreshold impedance profile fitted models than experimentally determined median electrotonic lengths of 0.78 for CA1 pyramidal neuron apical dendrites (Turner & Schwartzkroin, 1983). This makes our reduced models currently unsuitable for studying distal synaptic inputs that impinge more than 0.45 electrotonic lengths away from the soma. Further constraints on the postsynaptic potential (PSP) attenuation and impedance phase profile in our model would make these models useful for studying synaptic integration while keeping computational costs lower than for full-scale models with detailed morphology.

We predict that correcting for DBLO may lead to at least three substantial alterations to the behavior of neural network models: E and I synaptic summation, firing bistability, and neuromodulator sensitivity. We address these in turn.

### Implications for E/I balance and synaptic summation

Many real neural networks exhibit bursting activity in excitatory neurons, and modern models are able to capture these effects (Golomb et al., 2006; Roy & Narayanan, 2023; Tang et al., 2010). DBLO kicks in whenever the postsynaptic cell is firing briskly, such as in bursts (Moore et al., 2009). (Bianchi et al., 2022) showed in their computational study that low DBLO models attenuated theta range synaptic activity while high DBLO models amplified gamma range synaptic activity. The presence of a high baseline can more than double the effective weight of a typical GABA-ergic synapse due to the increase in driving force. If we consider the LJP-corrected chloride reversal potential to be around −87 mV (Atallah & Scanziani, 2009; Barmashenko et al., 2011), and if DBLO raises the mean potential from around −82 mV to around −65 mV, we have a more than 200% increase in GABA receptor’s driving force. Thus, the EI balance in models using neurons that lack DBLO, such as (Yu et al., 2014), may be impacted. This effect on inhibitory inputs is still more pronounced over brain development since the GABA receptor’s reversal potential starts close to resting potential (Owens & Kriegstein, 2002; Peerboom & Wierenga, 2021). While very few network models consider early neuronal physiology, it is clear that the outcome of GABA-ergic inputs may flip from none, or even excitatory, to inhibitory when high activity results in DBLO. The effect of DBLO on the synaptic summation of excitatory inputs is predicted to be smaller than that for inhibitory inputs since the driving potential shift is only about DBLO/(E_glu_-E_rest_) or about 30%. Nevertheless, this, too, is a substantial effect during burst activity.

### Implications for firing bistability

In Fig. 9, we showed that low DBLO models need very high levels of Na_P to exhibit bistable firing which may be unphysiological. Thus, this might make some low DBLO neurons resistant to Na_P-induced firing bistability. Although our highly schematized ball-and-stick models with firing bistability were based on CA1 pyramidal neuronal properties, their reduced morphology makes them reasonable proxies for other neuron subtypes that exhibit bistable firing. These include cortical pyramidal neurons of prefrontal cortex (Thuault et al., 2013), entorhinal layer III neurons (Tahvildari et al., 2007), subiculum neurons (Kawasaki et al., 1999), rostral ambiguus neurons (Rekling & Feldman, 1997), spinal motoneurons (Lee & Heckman, 1998), and anterior cingulate cortex pyramidal neurons (Zhang & Séguéla, 2010). Further analysis and more neuron-specific channel kinetic data will be required to show to what extent this constraint applies across different classes of neurons. In principle, the DBLO linkage to Na_P firing bistability has numerous implications for network function. For example, oscillations and epileptic activity have been linked to firing bistability (Fuentealba et al., 2005; Protachevicz et al., 2019). Bistability has also been proposed to expand the dynamic repertoire of network models, especially attractors for memory (Lundqvist et al., 2010).

### Implications for neuromodulation

Neuromodulators are ubiquitous control signals for causing networks to switch between states (Gutierrez & Marder, 2014; Marder et al., 2014). While they have been extensively analyzed in small CPGs (Golowasch, 2019; Williams et al., 2022), they also play a key role in setting network learning rates (Doya, 2002; Jepma et al., 2016; Rutledge et al., 2009). One of the key mechanisms by which neuromodulators affect neuronal activity is by modulating ion channels, which can be done by channel insertion or by phosphorylation (Nadim & Bucher, 2014). An accurate understanding of DBLO is relevant to this in two ways. First, many neuromodulated ion channels are potassium channels (Palacios-Filardo & Mellor, 2019). As discussed above for the case of inhibitory currents, the currents carried by potassium channels will be larger during episodes of DBLO than at resting potential, driven by an increase in their driving force. This means that DBLO will selectively amplify the effect of such neuromodulators at times of high activity. Second, the current study shows that models that correctly express high DBLO have significantly different channel densities than incorrect models (Fig. 9A). Thus, the substrate for neuromodulation itself (by way of density and distribution of K^+^ channels in particular) may be quite different. This has implications for studies that try to understand single-neuron activity changes arising from neuromodulation. Simply put, the correct DBLO model may respond differently to a neuromodulator than a model with similar firing activity but no DBLO.

## Materials and methods

All code and experimental data are freely available on GitHub at: https://github.com/BhallaLab/DBLO_paper. All simulations were carried out in MOOSE (Ray & Bhalla, 2008) except for Fig. 2, where NEURON was used to run the state-of-the-art models from ModelDB. These were performed on a 128-core AMD EPYC 7763 CPU machine with 1 TB RAM and running Ubuntu Server 22.04. For hypothesis testing, the Pingouin (Vallat, 2018), Statsmodel (Seabold & Perktold, 2010), and Scipy (Virtanen et al., 2020) Python libraries were used.

### Ion channel implementation

Eight ion channel types were used in this study, namely transient sodium channel (Na_T), persistent sodium channel (Na_P), slow delayed rectifier channel (K_DR), HCN channel (h), N-type calcium channel (CaN), L-type calcium channel (CaL), R-type calcium channel (CaR), and T-type calcium channel (CaT). Below, we describe their kinetics in brief.

HCN channel kinetics was taken from (Hay et al., 2011), which in turn were based on the recordings of apical dendrites of cortical layer 5 pyramidal neurons of adult male Wistar rats (Kole et al., 2008). E_rev_ of HCN channels was kept at −40 mV. The equations that describe the kinetics are as follows:

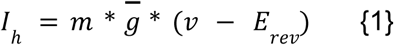

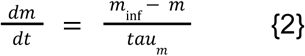

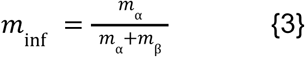

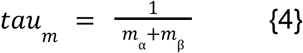

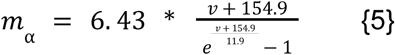

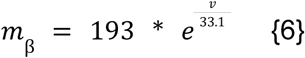

Where v is in mV.

Three formulations of Na_T kinetics from different papers have been used in this study. Unless otherwise stated, the steady state or the inf curve of Na_T channel kinetics was taken from experimental recordings by (Royeck et al., 2008) in the soma and AIS of mouse CA1 pyramidal neurons. Since (Royeck et al., 2008) did not have data about the time constants, those were taken from Hay et al., 2011. We chose not to use the kinetics used in (M. Migliore et al., 1999) as the default Na_T kinetics as it was based on dendritic recordings from CA1 pyramidal neurons of adult Sprague-Dawley rats, and the inactivation gate in the paper was modified. E_rev_ of Na_T channels was kept at 60 mV. The equations that describe the kinetics are as follows:

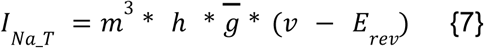

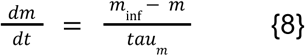

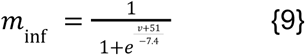

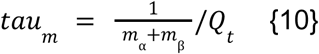

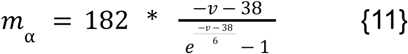

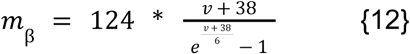

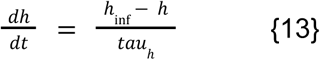

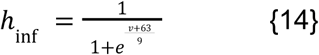

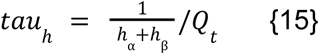

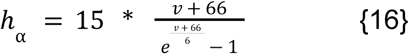

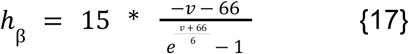

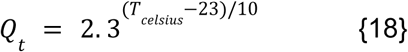

Where v is in mV.

The second Na_T kinetics formulation was from (R. Migliore et al., 2018). This was only used in parts of Fig. 8. The third Na_T kinetics formulation was from (Gouwens et al., 2018). This kinetics was used for parts of Fig. 8 and parts of Fig. 9.

The K_DR kinetics was taken from (R. Migliore et al., 2018) but it was parameterized and written in a different form as described below:

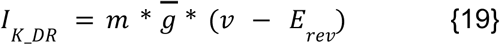

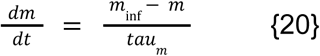

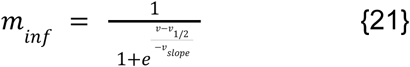

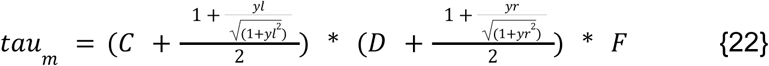

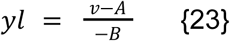

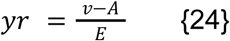

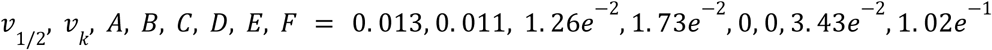

The parameterization was needed to explore the kinetics space that allows high DBLO. We tried different unimodal functions that can be used to parameterize the tau curves and found the eq{22-24} to be the most suitable. The A parameter controls the location of the peak of the tau curve, the F parameter controls the peak, the C parameter controls the tau at high V_m_, the D parameter controls the tau at low V_m_, the B parameter controls the slope of the tau at high V_m_, and the E parameters controls the slope of the tau at high V_m_ (see Fig. 7B for a visual representation). Its E_rev_ was kept at −100 mV unless otherwise stated.

The base Na_P channel kinetics was adapted from experimental recordings of pyramidal neurons in rat sensorimotor cortex (A. M. Brown et al., 1994). Briefly, it was modeled with a single activation gate with a time constant of 1 ms, V_1/2_ of −50.4 mV and k of 5.53 mV. This ion channel was only used for Fig. 9D. Its E_rev_ was kept at 60 mV.

The CaL, CaN, and CaT kinetics were taken from the CA1_PC_cAC_sig5 model of (R. Migliore et al., 2018). CaR kinetics, on the other hand, was taken from (Upchurch et al., 2022). The driving force of these channels was modeled with the Goldman-Hodgkin-Katz flux equation (Hille, 2001).

### State-of-art models

We used several CA1 pyramidal neuron models from ModelDB (McDougal et al., 2017): Migliore et al., 2018 (accession number: 244688); Turi et al., 2019 (accession number: 246546); and Tomko et al., 2021 (accession number: 266901). The (Gouwens et al., 2018) cortical pyramidal model was downloaded from http://celltypes.brain-map.org. The model ID was 486132712. For the Migliore 2018 model and the Turi 2018 model, where LJP was not corrected, we also made a version of the model where we modified the E_m_ such that the resting membrane potential of the model came down by 15 mV. In each of the models, a 500 ms current pulse of different amplitudes was injected. The amplitude was chosen such that the number of spikes in each model during the 500 ms current injection matches the 150 pA representative experimental recording shown in Fig. 1A.

### Simulations of activation and inactivation gates

To record the opening and closing of Na_T gates in simulations, we took a representative 300 pA experimental recording. We used the Na_T kinetics equations {8-18} to numerically calculate the activation and inactivation gate opening probability values at each time point. We did the same calculation for a 300 pA experimental recording that was not corrected for LJP.

### The AIS model

The AIS models were generated with two compartments. The somatic compartment had dimensions of 15 µm x 15 µm, while the AIS compartment had a length of 30µm and a diameter of 2 µm. E_m_ was fixed at −80 mV. Na_T and K_DR channels were included only in the AIS compartment. Random values of the free parameters, as listed in Table 4, were picked, and several models were generated. Their R_in_, C_in_, E_rest_, tau_m_ (membrane time constant), number of spikes in response to 500 ms of 0 pA current injection, number of spikes in response to 500 ms of 150 pA current injection, and average interspike interval (ISI) of APs in response to 500 ms of 150 pA current injection were calculated. If these features were not within the experimental range, the models were discarded. For subthreshold features, we considered the absolute experimental range, while for suprathreshold features, we considered the 10-90^th^ percentile of the experimental range. See Table 5 for subthreshold feature ranges and Table 6 for suprathreshold feature ranges. Out of 5000 models tested, we got a total of 457 valid models. We also calculated each model’s DBLO and spike height. The current injection and voltage recording, in this case, was done from the somatic compartment.

**Table 4:** Free parameters and their ranges for the unbiased stochastic search to test the AIS hypothesis. Parameters with * were not sampled uniformly but logarithmically with base 10.

| Free parameter (units) | Symbol | Lower limit | Upper limit |
| --- | --- | --- | --- |
| Specific membrane resistance ($\Omega m^2$) | RM | 0.01 | 0.2 |
| Specific axial resistance ($\Omega m$)* | RA | 1e-5 | 1e2 |
| Specific membrane capacitance (F/$m^2$) | CM | 0.086 | 0.124 |
| AIS Na<sub>T</sub> maximal conductance ($\mu S$)* | Na<sub>T</sub>_Gbar | 0.1 | 10 |
| AIS K<sub>DR</sub> maximal conductance ($\mu S$)* | K<sub>DR</sub>_Gbar | 1 | 1000 |

**Table 5:**
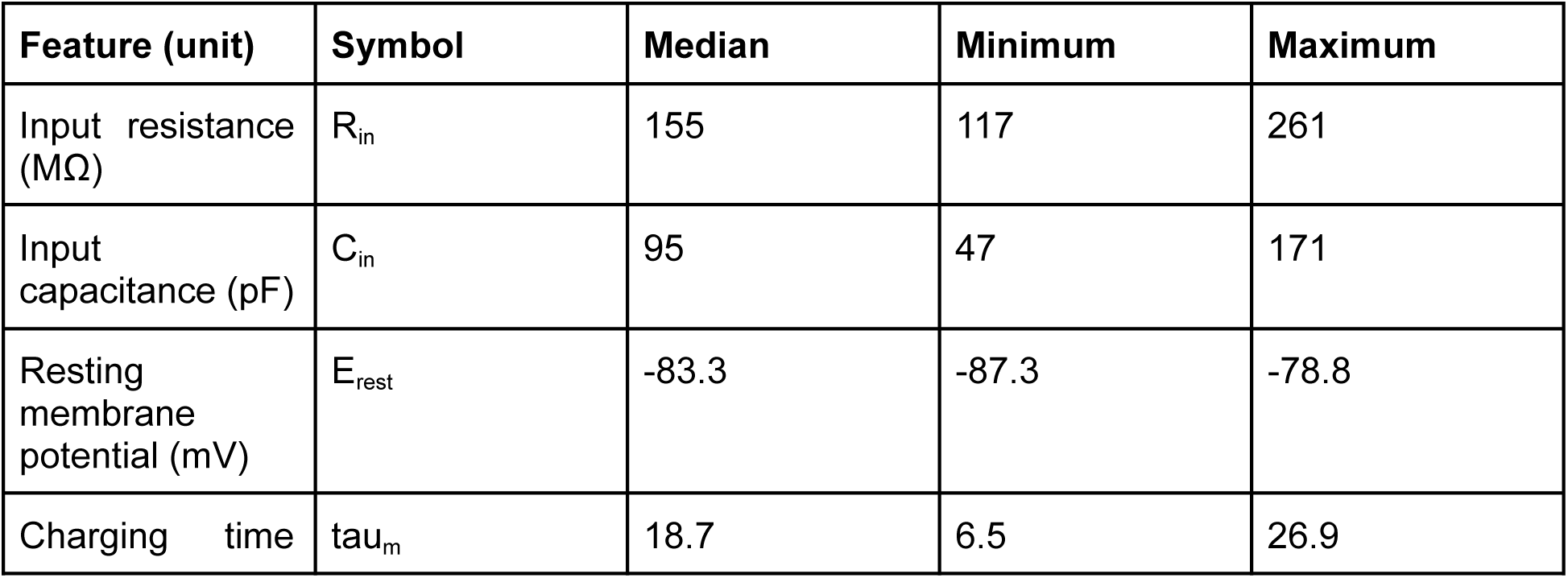
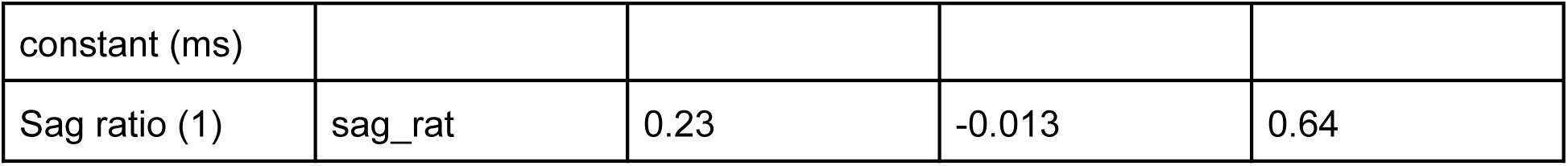
Subthreshold features of experimentally recorded neurons.

**Table 6:**
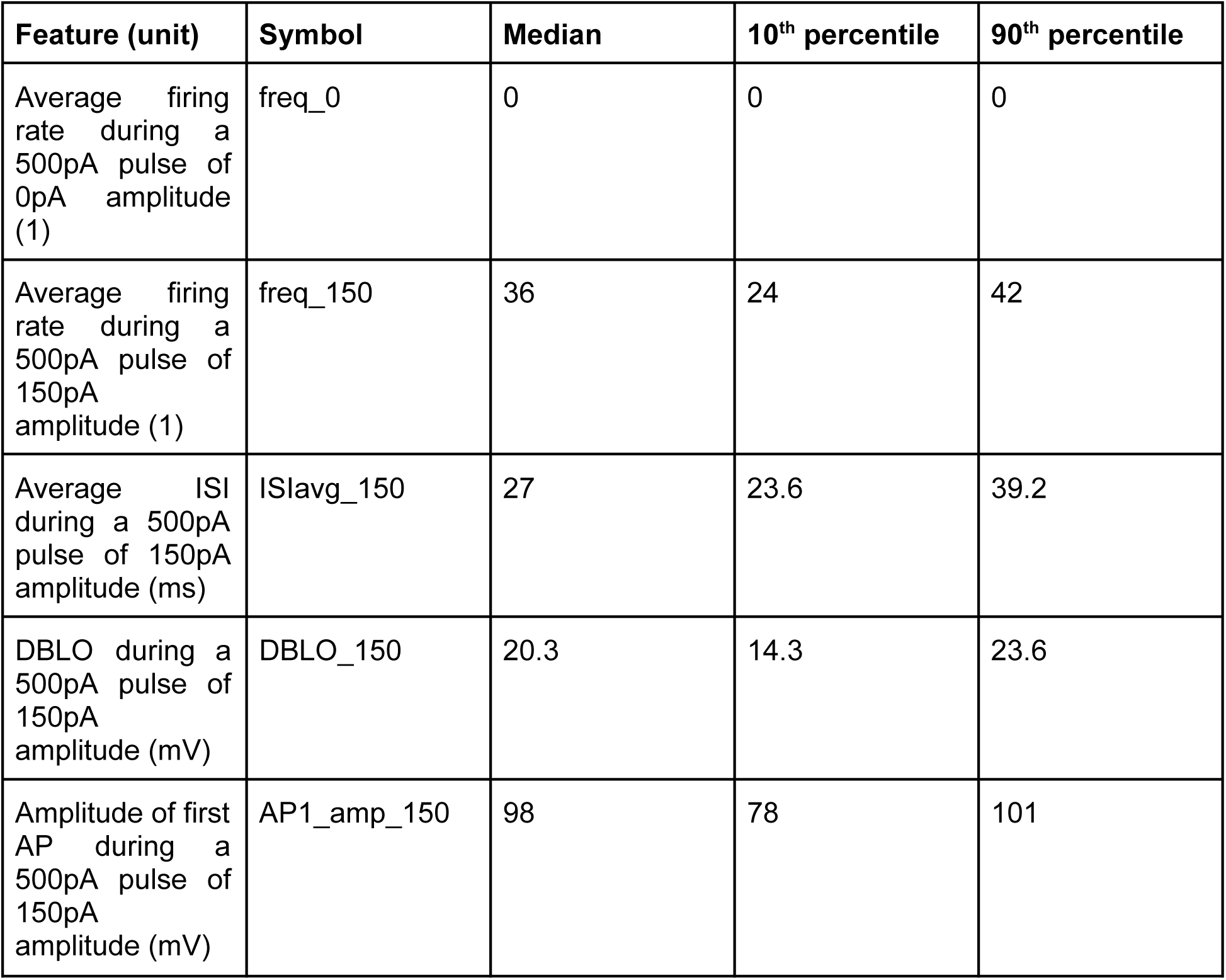
Suprathreshold features of experimentally recorded neurons.

### Animal care

All animals used for the experiments were BL6/J mice. They were raised with adequate water and food provided in cages with a cage strength of up to five. All animals underwent a 14-hour light and 10-hour dark cycle. Experimental animals were six to ten weeks old. Institute Animal Care and Resource Center (ACRC) staff ensured the health status, pathogen exposure, and well-being of all animals throughout the project timeline.

### Ethics

All protocols performed were approved by the NCBS Institutional Animal Ethics Committee. Approval details: NCBS-IAE-2016/ 21 (ME); NCBS-IAE-2016/ 7(M); NCBS-IAE-2022/8 (R2M), Animal facility registration number: 109/GO/ReRcBiBt/S/99/CPCSEA.

### Slice preparation

The slices were prepared from 6-10 weeks old (N=3) of BL6/J mice. Animals were decapitated after cervical dislocation, post-anesthesia with Halothane. 400 um thick slices were made from the dissected hippocampus on a vibratome (LEICA VT-1200). The dissections and slicing were performed in cutting solution, artificial cerebrospinal fluid (aCSF) with (in mM): NaCl-87, KCl-2.5, NaHCO3-25, NaH2PO4.H2O-1.25, Sucrose-75, MgCl2-7, CaCl2--0.5. Once sliced, the slices were transferred to a holding aCSF with (in mM): NaCl-124, KCl-2.7, NaHCO3-26, NaH2PO4.H2O-1.25, Glucose-10, MgCl2-1, CaCl2-2. Both solutions were saturated with 95% O2 and 5% CO2. After an incubation time of 45 mins, slices were transferred to the recording chamber with holding-aCSF perfusion at 31 ± 1 °C.

### Electrophysiology

We used an infrared Differential Interference Contrast (DIC) microscope(Olympus BX51WI) with 40x (Olympus LUMPLFLN, 40XW) water immersion objectives to visualize the cells. The electrophysiology data was acquired using Axon™ Digidata® 1440B digitizer and MultiClamp 700B amplifier (Molecular devices, Axon instruments). We used a 20 KHz sampling frequency and a 0-10KHz Bessel bandpass filter for the acquisition of electrical signals. We performed whole-cell patch recordings from CA1 pyramidal cells. The electrodes were made from thick-walled borosilicate glass capillaries (OD-1.5mm, ID-0.86mm) from Sutter Instruments, Novato, CA. We used a P-1000 pipette puller from Sutter Instruments, Novato, CA, with a box filament. Electrodes were filled with (mM): K-Gluconate-130, NaCl-5, HEPES-10, Na4-EGTA-1, MgCl2-2, Mg-ATP-2, Na-GTP-0.5, Phosphocreatine-10, pH adjusted to 7.3 and osmolarity adjusted to ∼290 mOsm. LJP was 15 mV. It was calculated according to the stationary Nernst–Planck equation (Marino et al., 2014) using LJPcalc software (https://swharden.com/LJPcalc). Experimentally recorded voltages were corrected for LJP except when stated otherwise.

Cells were picked for analysis based on the stability of series resistance (input resistance change less than 20% between the start of the recording and the end of the recording) and resting membrane potential (a shift of less than 2.5 mV between the beginning of the recording and the end of the recording). Out of 14 cells that we recorded, 13 cells passed the criterion.

### Recording chirp responses from mouse CA1 pyramidal neurons

To record the subthreshold impedance amplitude of neurons, we injected a chirp stimulus with time-varying frequency into the neuron. We recorded its voltage response using a whole-cell clamp setup (as described above). The chirp stimulus was generated using the function:

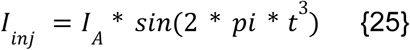

Where I_A_ is 30 pA

We injected this current for 13s. During this time, the frequency went from 0 Hz to 500 Hz in a quadratic manner (Fig. 6B).

The impedance amplitude was then calculated as:

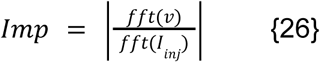

### Subthreshold feature calculation

To measure the subthreshold features of both experimentally recorded neurons and models, a 500 ms pulse of −25 pA current amplitude was injected. The first 100 ms of the −25 pA current injection response was fitted to the following equation:

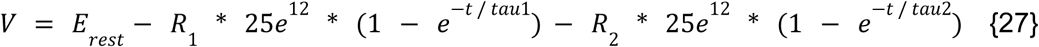

The R_in_ (input resistance) was defined as the higher of the two parameters, R_1_ and R_2_. The higher of the two parameters, tau1 and tau2, was taken as tau_m_ (membrane time constant), and C_in_ (input capacitance) was calculated as tau_m_/R_in_.

The E_rest_ was calculated as the median membrane potential during 0 pA current injection of 500 ms in both models and experimental recordings.

The steady-state voltage response (V_ss_) at −25 pA current injection was calculated as the median membrane potential of the last 100 ms of the voltage response to the 500 ms pulse.

The subthreshold features, along with their median and experimental range, are given in Table 5.

### Suprathreshold electrophysiological feature calculation

To calculate the suprathreshold electrophysiological features, we used a combination of BlueBrain’s eFEL library (Ranjan et al., 2024) and custom code. Table 6 summarizes these features for the experimentally recorded neurons.

The average ISI feature was needed when generating valid active models to reject models with a valid number of APs that exhibited irregular firing.

### Generating valid subthreshold single-compartment models

To generate valid subthreshold single-compartment models (termed *1compt* models), we used a single cylindrical compartment with a diameter of 15 µm and length of 15 µm. We employed a custom algorithm (included in the GitHub repository of this paper) to generate valid models. The algorithm combines stochastic search and gradient descent to generate a single valid subthreshold model corresponding to each of the recorded CA1 pyramidal neurons. In the first step of the stochastic search, we picked uniformly distributed random values for three free parameters, namely - membrane resistance of the compartment (sm_RM), total capacitance of the compartment (sm_CM), and maximal conductance of the HCN channel (h_Chan_Gbar). E_m_ was then calculated to give the same E_rest_ as the corresponding neuron’s E_rest_. Table 7 shows the range from which these values were picked. We generated 1000 such random models and recorded their response to 500ms of −25 pA current injection (V_-25_). Then, for each of the experimentally recorded cells, we found the 10 models out of 1000 that had the least root mean square error (RMS) between the V_-25_ of the model and the experimentally recorded cell. Thus, we got the 10 best models corresponding to each neuron. We then employed Scipy’s curve_fit function to fit these 10 best models to the corresponding neuron’s V_-25_. Finally, we picked the best out of the 10 to get a single valid passive single-compartment model for each of the 13 experimentally recorded neurons for a total of 13 models.

**Table 7:**
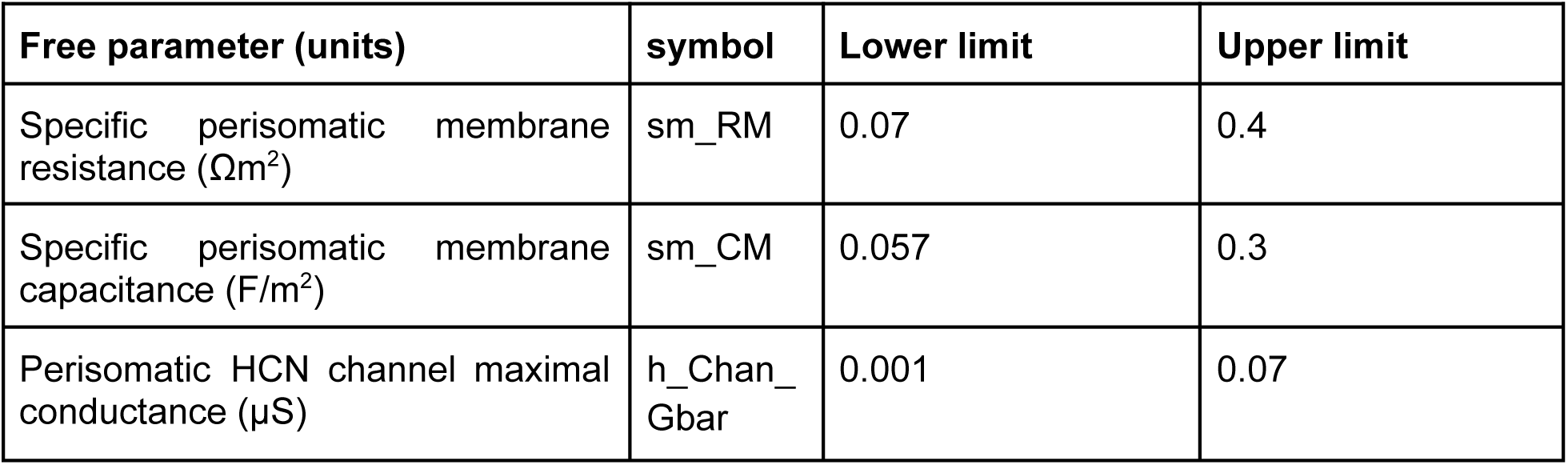
Free parameters and their ranges for the unbiased stochastic search to generate valid subthreshold *1compt* models.

### Generating valid subthreshold features fitted ball and stick models

To generate valid passive ball and stick models (termed *pas* models), we used a morphology with a perisomatic compartment and a single dendrite connected to the perisomatic compartment. The perisomatic compartment had a diameter of 15 µm and a length of 15 µm. These dimensions were chosen to be similar to the somatic dimensions of mouse CA1 pyramidal neurons. The dendritic section had a diameter of 4 µm and a length of 500 µm. It was divided into 10 compartments (referred to as segments in NEURON nomenclature) to ensure that each compartment was small enough to follow the d_λ_ rule. The d_λ_ rule states that each segment is at most 0.1 times the AC length constant at 100 Hz (Hines & Carnevale, 2001). To account for sag and inductance in the subthreshold response of a neuron, we included an HCN channel in the perisomatic compartment. We employed a similar approach to the one described in the last section, except with a few modifications. There were 6 free parameters instead of 4, namely - perisomatic compartment RM, perisomatic compartment CM, perisomatic compartment axial resistance (RA), dendritic compartment RM, dendritic compartment CM, and dendritic compartment RA (table 8). The E_m_ and h_Chan_Gbar for the models were taken from the previously generated *1compt* models. Instead of using V_-25_ for fitting, we used the corresponding neuron’s R_in_ and tau_m_. Also, unlike for the *1compt* case, we generated 10 valid models for each neuron (for a total of 130 models) instead of just one valid model due to the degeneracy that exists in this case. Many passive parameter combinations can give rise to the same subthreshold features while having considerable differences in their subthreshold impedance profile.

**Table 8:** Free parameters and their ranges for the unbiased stochastic search to generate valid passive ball and stick models and valid impedance amplitude fitted models.

| Free parameter (units) | symbol | Lower limit | Upper limit |
| --- | --- | --- | --- |
| Perisomatic membrane resistance ($\Omega m^2$) | sm_RM | 0 | 20 |
| Perisomatic membrane capacitance (F/$m^2$) | sm_CM | 0 | 0.2 |
| Perisomatic axial resistance ($\Omega m$) | sm_RA | 0 | 20 |
| Dendritic membrane resistance ($\Omega m^2$) | dend_RM | 0 | 20 |
| Dendritic membrane capacitance (F/$m^2$) | dend_CM | 0 | 0.2 |
| Dendritic axial resistance ($\Omega m$) | dend_RA | 0 | 20 |

### Generating valid subthreshold impedance amplitude profiles matched ball and stick models

The same morphology and fitting procedure as described above were used, except the models were fitted to the subthreshold impedance amplitude profile of each of the 13 cells. The resulting models fitted well to both the −25 pA response and the subthreshold impedance amplitude profiles of the corresponding neuron. For this case, too, we generated a single valid model corresponding to each neuron.

### Generating valid active model triplets

For the morphology hypothesis, we generated triplets of valid active models. For each neuron, we picked its corresponding *1compt* subthreshold model, *imp* model, and one of the ten *pas* models we generated earlier. We then uniformly sampled Na_T_Gbar and K_DR_Gbar over the range provided in Table 9 and incorporated these Gbars into the triplet of subthreshold models. We then checked whether any of these models give suprathreshold features outside the experimental range for freq_0, freq_150, ISIavg_150, and AP1_amp_150 (see Table 6 for the experimental range). If any, the whole triplet was discarded. We repeated this procedure for a neuron 5000 times. Thus, we inspected a total of 13×5000=65000 triplets. A total of 4174 triplets were found to be valid and were used for analysis in Fig. 5. This procedure allowed us to compare models with the same maximal conductances and subthreshold features but different impedance amplitude profiles.

**Table 9:** Free parameters and their ranges for the unbiased stochastic search to generate active model triplets (*1compt*, *pas*, and *imp*). *The Gbars were logarithmically sampled with a base of 10.

| Free parameter (units) | symbol | Lower limit | Upper limit |
| --- | --- | --- | --- |
| Perisomatic Na<sub>T</sub> channel maximal conductance (μS)* | Na_T_Chan_Gbar | 0.1 | 10 |
| Perisomatic K<sub>DR</sub> channel maximal conductance (μS)* | K_DR_Chan_Gbar | 1 | 1000 |

### Generating models with different K_DR reversal potential

Single-compartment models of dimensions 15 µm (length) x 15 µm (diameter) were used. The free parameters and their allowed range are given in Table 10. We used a uniform random distribution to pick the values of these free parameters and recorded their R_in_, C_in_, E_rest_, freq_0, freq_150, and ISIavg_150. Since we were modifying the K_DR E_rev_ to unphysiologically high levels, the validity criteria were a bit relaxed for freq_150 and ISIavg_150. Instead of using the 10-90^th^ percentile experimental range as the valid range, we used half the 10^th^ percentile experimental value and double the 90^th^ percentile experimental value as the valid range (see Table 6 for the 10^th^ and 90^th^ percentile experimental values of the active features). For subthreshold features, the validity criterion was still the same (table 5). If these values were not within the experimental range, the model was discarded. We also recorded each valid model’s DBLO at 150 pA current injection. We did not include h_Chan in these models. Out of a total of 5000 models that were inspected, 231 were valid and were used for analysis.

**Table 10:**
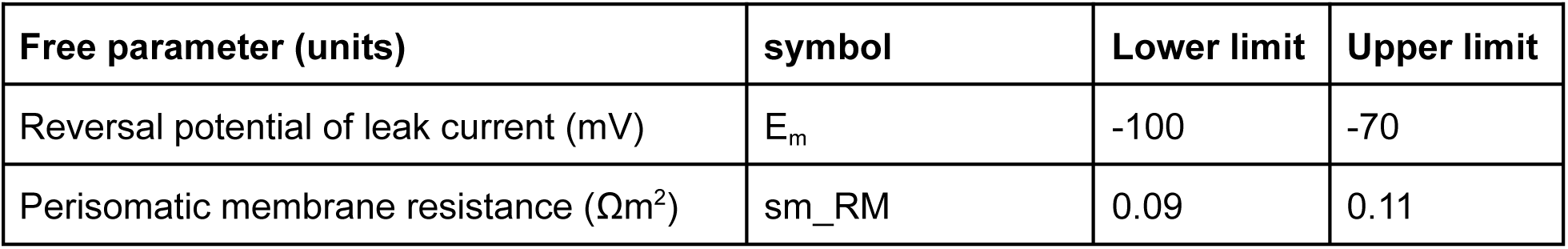
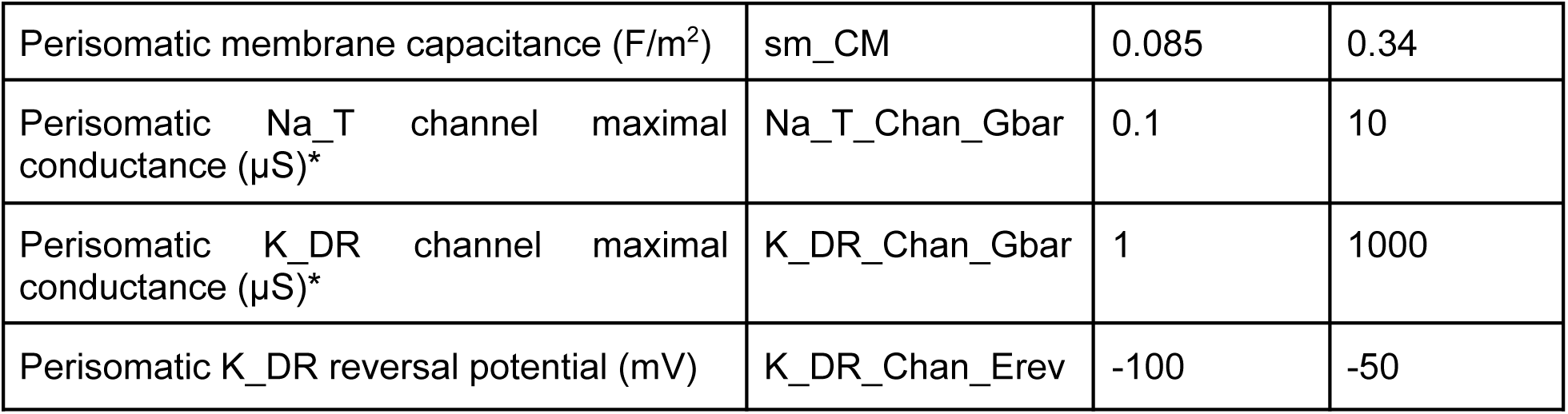
Free parameters and their ranges for the unbiased stochastic search to generate active models for the K_DR E_rev_ hypothesis. Parameters with * were not sampled uniformly but logarithmically with base 10.

### Exploring K_DR kinetics space

To build models with different K_DR kinetics, we randomly picked one of the thirteen subthreshold *1compt* models we built previously and incorporated Na_T_Chan and K_DR_Chan in those models. The free parameters are provided in Table 11. Out of the eight free parameters, six were for the K_DR time constant curve parameterized as in eq {22-24}. These parameters are further explained in Fig 7B and eq {22-24}. We did not vary the steady state inf curve of K_DR. A random number with a uniform distribution between the limits specified in the table was picked for each parameter. The model was then checked to see whether their freq_150, freq_0, ISIavg_150, and AP1_amp_150 were within the experimental range (Table 6). This whole procedure was repeated a total of 50000 times, and the generated models were checked for validity. 677 models passed the validity criterion.

**Table 11:**
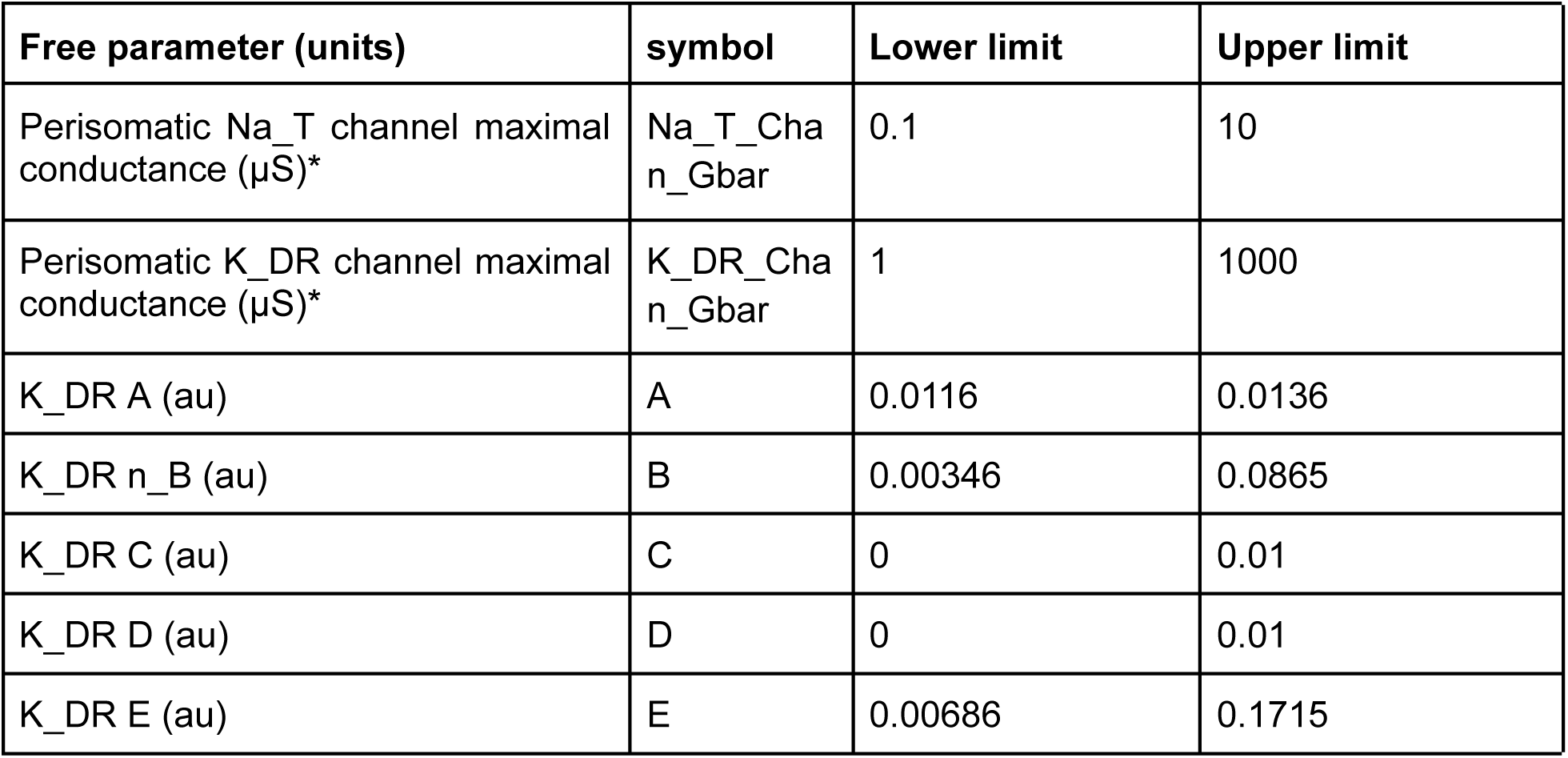
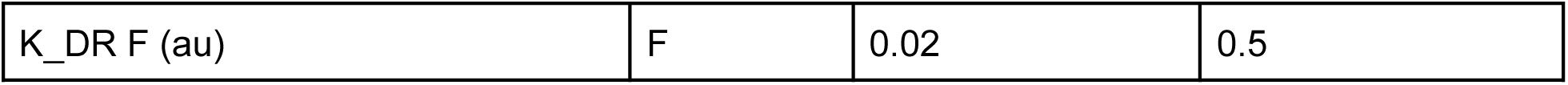
Free parameters and their ranges for the unbiased stochastic search to explore the K_DR kinetics space.

### Models with different Na_T kinetics

For this part, we made three families of models - *Gou*, *Mig*, and *Roy*. For each of these families of models, we first picked one of the 13 subthreshold *1compt* models. We then incorporated Na_T and K_DR into those models with their Gbar levels chosen uniformly over the range specified in Table 12. For *Gou* models, Na_T kinetics from Gouwens et al., 2018 was used as described in a previous section. For the *Mig* models, the Na_T kinetics were derived from (R. Migliore et al., 2018). For the *Roy* and *Gou* models, they were derived from (Royeck et al., 2008) and (Hay et al., 2011), respectively. Refer to the first section of Materials and Methods for details on the kinetics. The models were then checked to see if their freq_0, freq_0, and AP1_amp_150, and ISIavg_150 were within the experimental range (table 6). Invalid models were discarded. This procedure was repeated 500 times for each subthreshold *1compt* model for a total of 13×500=6500 times per model family. For *Gou* models, 406 models were valid. 119 *Roy* models were valid, while only 24 *Mig* models were valid.

**Table 12:**
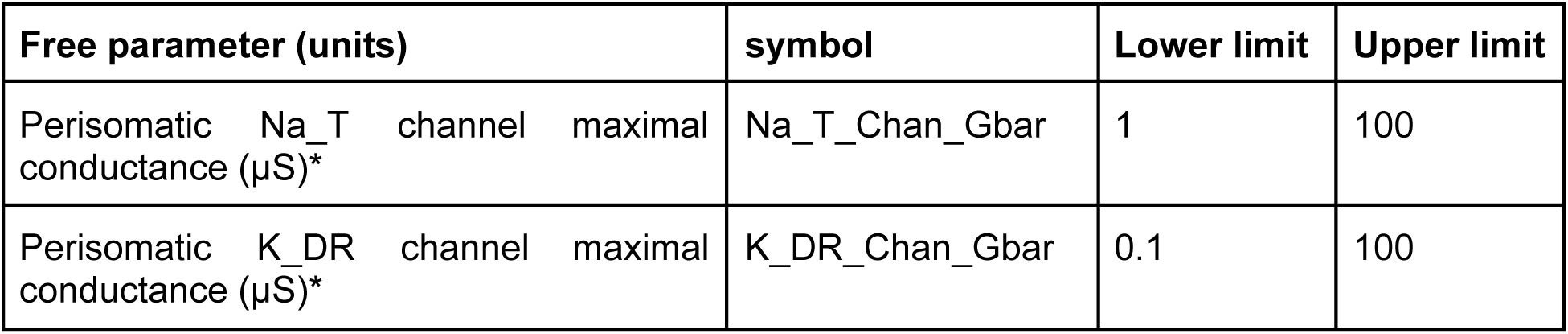
Free parameters and their ranges for the unbiased stochastic search to generate models with different Na_T kinetics (*Gou*, *Roy*, and *Mig*). *The Gbars were logarithmically sampled with a base of 10.

### Generating valid unified models

To generate valid consolidated models, we started by randomly picking one of the thirteen subthreshold *imp* models. We incorporated Gouwens et al., 2018 Na_T kinetics and a fast deactivating K_DR channel in the model. The fast deactivating K_DR kinetics was generated by dividing the E parameter by 2 in the parameterized original (R. Migliore et al., 2018) K_DR kinetics. To compensate for the change in the resultant curve’s peak, the A parameter was shifted right by 0.006, and the F parameter was multiplied by 1.1 (see eq {22-24} for the kinetics formulation). A depolarized K_DR E_rev_ of −90mV was used in this case. The Gbar values were chosen from a uniform distribution with the range specified in Table 13. We then checked whether the model’s freq_0, freq_150, ISIavg_150, and AP1_amp_150 were within the experimental range (table 6). Invalid models were discarded. We repeated the above procedure 5000 times for each subthreshold *imp* model for a total of 13×5000=65000 models. 1171 models passed this validity test.

**Table 13:** Free parameters and their ranges for the unbiased stochastic search to generate valid unified models. *The Gbars were logarithmically sampled with a base of 10.

| Free parameter (units) | symbol | Lower limit | Upper limit |
| --- | --- | --- | --- |
| Perisomatic Na<sub>T</sub> channel maximal conductance (μS)* | Na<sub>T</sub>_Chan_Gbar | 1 | 100 |
| Perisomatic K<sub>DR</sub> channel maximal conductance (μS)* | K<sub>DR</sub>_Chan_Gbar | 0.1 | 100 |

### Linear mixed-effect model (LMM)

In order to assess the correlation between Na_T_Chan_Gbar/K_DR_Chan_Gbar and DBLO levels in our valid unified models, we used an LMM as implemented in the statsmodels package (statsmodels.formula.api.smf.mixedlm). Given that the unified models were derived from 13 base subthreshold *imp* models (see previous section), we accounted for variability across these base models as random effects while treating DBLO as a fixed effect. The intercepts were allowed to vary between groups, while the slope of DBLO was modeled across all groups as a fixed effect. The LMM was fit using restricted maximum likelihood (REML) estimation.

### Models with calcium channels

We built four families of models, each with a different type of Calcium channel. For CaL models, in each of the 1171 valid unified models, we inserted the L-type calcium channel whose kinetics and maximal conductance values were taken from (R. Migliore et al., 2018). Similarly, for CaN, CaR, and CaT models. As (R. Migliore et al., 2018) did not use R-type calcium channels, we used the kinetics and maximal conductance from (Upchurch et al., 2022). After correcting for the soma area, the maximal conductance used for L-type, N-type, R-type, and T-type was 0.080 nS, 0.038 nS, 0.32 nS, and 0.0081 nS, respectively. For calcium dynamics, we used the following equation:

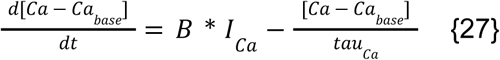

where Ca_base_ = 100 nM, tau_Ca_ = 80 ms, and B = 1.8e9 mM/A. These parameters were taken from (Gouwens et al., 2018) after standardizing for the soma area.

A 150 pA current was injected into the soma of all these models, and the internal calcium concentration was recorded during the AP train.

Note that the value of DBLO at 300 pA was calculated in the absence of any calcium channel.

### Bistable models

We first picked one of the 1171 active valid *unified* models that were generated previously. We incorporated Na_P channel into the model with kinetics as described previously. We then used a binary search to find the minimum Na_P_Chan_Gbar that would make the model show firing bistability. To check for firing bistability, we injected 150 pA to switch the model’s state from silent to firing. The bistable models maintained this state even after the 150 pA current injection phase. These models further switched to silent state on injection of −50 pA current. They continued to be silent after the −50 pA phase. We repeated the procedure for all the models. Out of 1171 models, 105 models did not show firing bistability even at very high Na_P_Chan_Gbar. We did not re-calculate DBLO or any other electrophysiological properties of these bistable models.

### Models with gap junctions

We built two instances of every valid unified model, which we named Neuron 1 and Neuron 2. We connected the soma of Neuron 1 and Neuron 2 by a gap junction. We injected 300 pA into the soma of Neuron 1 and recorded the APs from the soma of both instances. We then calculated, using a binary search algorithm, the minimum gap junction conductance needed for the 1^st^ AP generated in Neuron 1 to spread to the soma of Neuron 2. We performed the same analysis for the 2^nd^ AP. Note that the value of DBLO at 300 pA was calculated in the absence of any gap junction.

## Acknowledgments

NCBS-TIFR receives support from the Department of Atomic Energy, Government of India, under Project Identification No. RTI 4006. We thank Sufyan Ashhad for his valuable suggestions, especially regarding the role of K_DR’s reversal potential. We also thank Rishikesh Narayanan for his useful suggestions on the project.

## Author contributions

AK performed the simulations and wrote the paper. AKS performed the experiments and wrote the corresponding methods section. USB supervised the project, obtained funds, and wrote the paper.

## Data and Model availability

All data, models, and code used in this work are available on the GitHub repository: https://github.com/BhallaLab/DBLO_paper. The models are also available on ModelDB with accession number 2018017.

## Glossary

AHP: Afterhyperpolarization

AIS: Axon Initial Segment

AP: Action Potential

CM: Specific capacitance

DBL: Depolarization Baseline

DBLO: Depolarization Baseline Offset

E_glu_: Reversal potential of Glutamatergic ion channels

E_K_: Reversal potential of K^+^ channels

E_Na_: Reversal potential of Na^+^ channels

E_rest_: Resting membrane potential

E_rev_: Reversal potential

Gbar: Maximal conductance

HCN: hyperpolarization-activated

cAMP: dependent channel

HH: Hodgkin-Huxley

K_DR: Delayed rectifier potassium channel

Na_P: Persistent sodium channel

Na_T: Transient sodium channel

RA: Axial resistance

Rin: Input resistance

RM: Specific membrane resistance

SA: Sino-Atrial

Tau: Time constant

---

**Supplementary Fig. S1.**
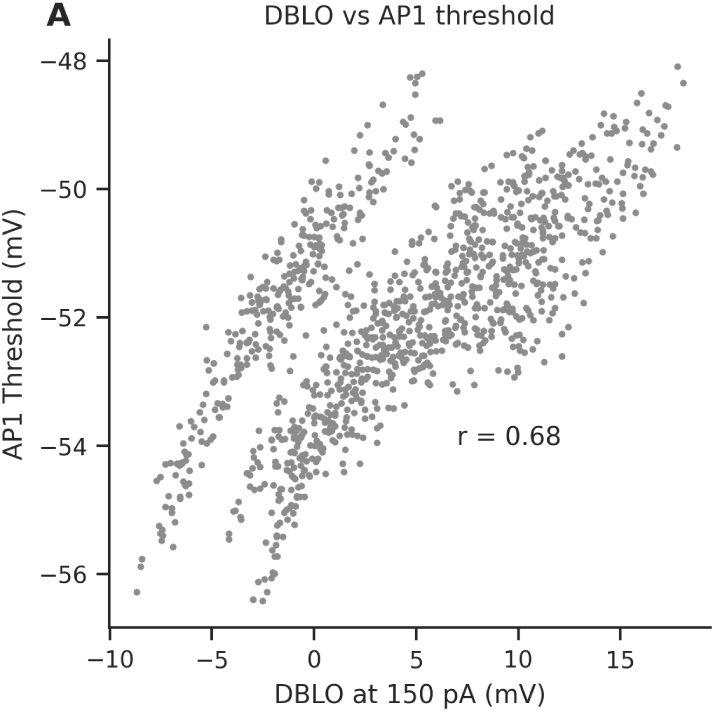
(A) DBLO is positively correlated with AP1 threshold. Pearson’s correlation coefficient = 0.68.

